# Single-cell growth inference of *Corynebacterium glutamicum* reveals asymptotically linear growth

**DOI:** 10.1101/2020.05.25.115055

**Authors:** Joris Messelink, Fabian Meyer, Marc Bramkamp, Chase P. Broedersz

**Author notes:** These authors contributed equally (Joris Messelink, Fabian Meyer). Corresponding authors: Chase Broedersz, Marc Bramkamp.

## Abstract

Regulation of growth and cell size is crucial for the optimization of bacterial cellular function. So far, single bacterial cells have been found to grow exponentially, which implies the need for tight regulation to maintain cell size homeostasis. Here, we characterize the growth behavior of the apically growing bacterium *Corynebacterium glutamicum* using a novel broadly applicable inference method for single-cell growth dynamics. Using this approach, we find that *C. glutamicum* exhibits asymptotically linear single-cell growth. To explain this growth mode, we model elongation as being rate-limited by the apical growth mechanism. Our model accurately reproduces the inferred cell growth dynamics and is validated with elongation measurements on a transglycosylase deficient *ΔrodA* mutant. Finally, with simulations we show that the distribution of cell lengths is narrower for linear than exponential growth, suggesting that this asymptotically linear growth mode can act as a substitute for tight division length and division symmetry regulation.

## Introduction

Regulated single-cell growth is crucial for the survival of a bacterial population. At the population level, fundamental laws of growth were revealed by Monod and Schaechter in the middle of the 20^th^ century, such as the identification of distinct population growth phases (Monod 1949; Schaechter, Maaløe, and Kjeldgaard 1958). However, at the time growth behavior at the single-cell level remained elusive. This changed over the last decade, as evolving technologies enabled detailed measurements of single-cell growth dynamics. Extensive work was done on common model organisms, including *Escherichia coli, Bacillus subtilis*, and *Caulobacter crescentus*, revealing that for these species single cells grow exponentially (Taheri-Araghi et al. 2015; Mir et al. 2011; Iyer-Biswas et al. 2014; Yu et al. 2017; Godin et al. 2010).

Such exponential growth is indeed expected if cellular volume production is proportional to the protein content (Amir 2014), as shown to be the case for *E. coli* (Belliveau et al. 2020). Importantly however, such a proportionality will only be present if cellular volume production is rate-limiting for growth. Cells with different rate-limiting steps could display distinct growth behavior, but little evidence exists for such deviations from exponential growth (Santi et al. 2013; Priestman et al. 2017). A promising candidate for uncovering novel growth modes is the Gram-positive *Corynebacterium glutamicum*. This rod-shaped bacterium grows its cell wall exclusively at the cell poles, allowing, in principle, for deviations from exponential single-cell growth (Figure 1). The dominant growth mode depends on the rate-limiting step for growth, which is presently unknown for this bacterium. Non-exponential growth modes may have important implications for growth regulatory mechanisms: while exponential growth requires checkpoints and regulatory systems to maintain a constant size distribution (Mir et al. 2011), such tight regulation might not be needed for other growth modes.

**Figure 1.**
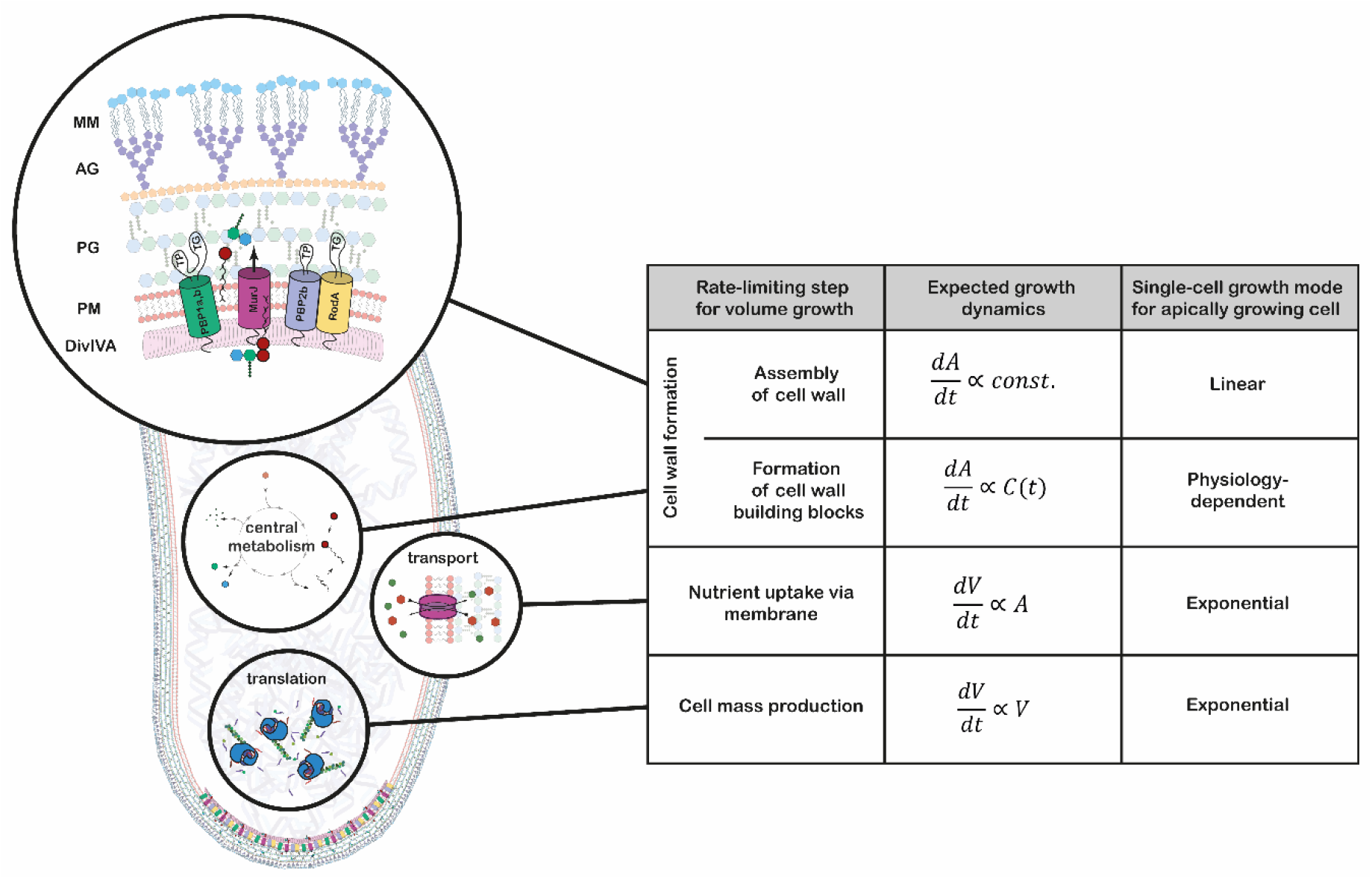
Growth mode analysis for four possible rate-limiting steps for cellular volume growth in the apically growing *C. glutamicum*. Here, *V* is the cellular volume, *A* is the cell wall area, and *C*(*t*) is the concentration of membrane building blocks in the cytoplasm. A constant cell width is assumed throughout, implying 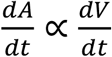. A fixed production capacity per unit volume is assumed for the rate-limiting steps ‘cell mass production’ and ‘formation of cell wall building blocks’. For the rate-limiting step ‘assembly of cell wall’, a constant insertion area at the cell poles is assumed. For an analysis of the single-cell growth mode if cell wall building block formation is the rate-limiting step for growth, see Appendix 1. Cell mass production, specifically ribosome synthesis, has previously been indicated as the rate-limiting step for growth in *E. coli* (Belliveau et al. 2020; Scott et al. 2010; Amir 2014). Linear growth is observed if the rate-limiting step for volume growth is the cell wall assembly (shown here in a simplified representation). The protein DivIVA serves as a scaffold at the curved membrane of the cell pole for the recruitment of the Lipid-II flippase MurJ and several mono- and bi-functional trans-peptidases (TP) and -gylcosylases (TG). In the process of elongation, peptidoglycan (PG) precursors are integrated into the existing PG sacculus, which serves as a scaffold of the synthesis of the arabinogalactan-layer (AG) and the mycolic-acid bilayer (MM).

*Corynebacterium glutamicum* is broadly used as a production-organism for amino-acids and vitamins and also serves as model organism for the taxonomically related human pathogens *Corynebacterium diphteriae* and *Mycobacterium tuberculosis* (Hermann 2003; Antoine, Coene, and Cocito 1988; Schubert et al. 2017). A common feature of Corynebacteria and Mycobacteria is the existence of a complex cell envelope. The cell wall of these bacteria is a polymer assembly composed of a classical bacterial peptidoglycan (PG) sacculus that is covalently bound to an arabinogalactan (AG) layer (Alderwick et al. 2015). Mycolic acids are fused to the arabinose and form an outer membrane like bilayer, rendering the cell surface highly hydrophobic (Puech et al. 2001). The mycolic acid membrane (MM) is an efficient barrier that protects the cells from many conventional antibiotics.

*C. glutamicum*’s growth and division behavior is vastly different to that of classical model species. In contrast to rod-shaped firmicutes and γ-proteobacteria, where cell-wall synthesis is dependent on the laterally acting MreB, members of the *Corynebacterianeae* lack a *mreB* homologue and elongate apically. This apical elongation is mediated by the protein DivIVA, which accumulates at the cell poles and serves as a scaffold for the organization of the elongasome complex (Letek et al. 2008; Hett and Rubin 2008; Sieger et al. 2013) (Figure 1, 2A, B). Furthermore, a tightly regulated division-site selection mechanism is absent in this species. Without harboring any known functional homologues of the Min- and nucleoid occlusion (Noc) system, division typically results in unequally sized daughter cells (Donovan et al. 2013; Donovan and Bramkamp 2014). Lastly, the spread in growth times between birth and division is much wider than in other model organisms, suggesting a weaker regulation of this growth feature (Donovan et al. 2013). These atypical growth properties suggest that this bacterium is an interesting candidate to test the universality of previously reported exponential growth laws. To reveal the underlying growth regulation mechanisms, it is necessary to study the elongation dynamics of *C. glutamicum*.

**Figure 2.**
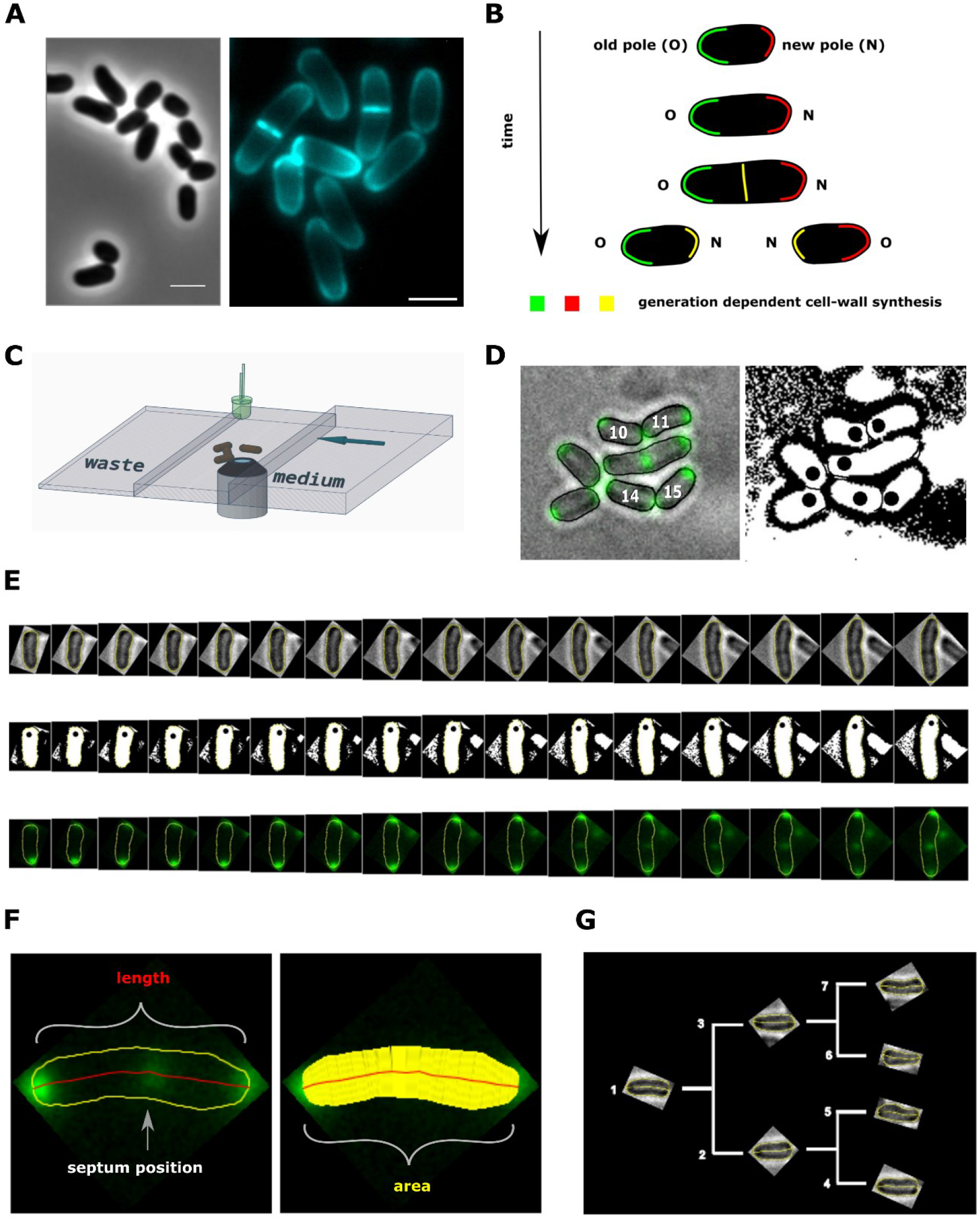
Experimental procedure and image analysis. **(A, left)** Phase contrast image of *C. glutamicum* in logarithmic growth phase, indicating the variable size of daughter cells. **(A, right)** HADA labeling of nascent peptidoglycan (PG), indicating the asymmetric apical growth where the old cell-pole always shows a larger area covered compared to the new pole. The labeling also reveals the variable septum positioning; Scale bar: 2 μm **(B)** Schematic showing the generation dependent sites of PG synthesis in *C. glutamicum*, including the maturation of a new to an old cell-pole. **(C)** Illustration of the microfluidic device for microscopic monitoring of a growing colony. **(E)** Example screen-shot of the developed method to extract individual cell cycles from a multi-channel time-lapse micrograph. The left panel shows a merging of the bright-field channel and the mCherry-tagged DivIVA together with an individual ID# that is assigned to cells right after division. The black dots in the right panel indicate the new cell pole. **(E)** Example of an extracted individual cell cycle from birth (left) until prior to division (right), showing the bright-field (top), the orientation (middle) and the localization of mCherry-tagged DivIVA (bottom). **(F)** Example of the developed single cell analysis algorithm, measuring the length according to the cell’s geometry, as well as the cell’s area and the septum position relative to the new pole. **(G)** Dendrogram providing the rationale for identification of single cells in a growing colony.

Here, we measure the single-cell elongations within a proliferating population of *C. glutamicum* cells, and develop an analysis procedure to infer their growth behavior. We find that *C. glutamicum* deviates from the generally assumed single-cell exponential growth law. Instead, these *Corynebacteria* exhibit asymptotically linear growth. We develop a mechanistic model, termed the rate-limiting apical growth (RAG) model, showing that this anomalous elongation dynamics is consistent with the polar cell wall synthesis being the rate-limiting step for growth. Finally, we demonstrate a connection between mode of growth and the impact of single-cell variability on the cell size distribution of the population. For an asymptotically linear grower, these variations have a much smaller impact on this distribution than they would for an exponential grower, suggesting an evolutionary explanation for the lack of tight regulation of single-cell growth in *C. glutamicum*.

## Results

### Measuring elongation trajectories using microfluidic experiments

To measure the development of single *C. glutamicum* cells over time, we established a workflow combining single-cell epifluorescence microscopy with semi-automatic image processing. Cells were grown in a microfluidic device. We used wild type cells and cells expressing the scaffold protein DivIVA as a translational fusion to mCherry. DivIVA is used as a marker for cell cycle progression, since it localizes to the cell poles and to the newly formed division septum in *C. glutamicum* (Letek et al. 2008; Donovan et al. 2013).

For the choice of microfluidic device, we deviate from the commonly used Mother Machine (Long et al. 2013), which grows bacteria in thin channels roughly equaling the cell width. The Mother Machine is not ideally suited for *C. glutamicum* growth, as the characteristic V-snapping at division could lead to shear forces and stress during cell separation, affecting growth (Bertozzi et al. 2019). Indeed, in some cases the mother machine has been shown to affect growth properties even in cells not exhibiting V-snapping at division, due to mechanical stresses inducing cell deformation (Yang et al. 2018). Therefore, we instead used microfluidic chambers that allow the growing colony to expand without spatial limitations into two dimensions for several generations (Figure 2C, D, Materials and Methods). Within the highly controlled environment of the microfluidic device, a steady medium feed and a constant temperature of 30 °C was maintained. We extracted bright-field- and fluorescent-images over three-minute intervals, which were subsequently processed semi-automatically with a workflow developed in FIJI and R (Schindelin et al. 2012; R Development Core Team 2003). For each individual cell per time-frame, the data set contains the cell’s length, area and estimated volume, the DivIVA-mCherry intensity profile, and information about generational lineage (Figure 2E-G). We used these data sets to further investigate the growth behavior of our bacterium. Thus, using this procedure, we obtained data sets containing detailed statistics on single-cell growth of *C. glutamicum*.

For subsequent analysis, the measured cell lengths were used, because of their low noise levels as compared to other measures (Appendix 2-Figure 1B). Importantly, the increases in cell length are proportional to the increases in cell area (Appendix 2-Figure 1A), suggesting that cellular length increase is also proportional to the volume increase. This proportionality is expected since the rod-shaped *C. glutamicum* cells insert new cell wall material exclusively at the poles, while maintaining a roughly constant cell width over the cell cycle (Schubert et al. 2017; Daniel and Errington 2003).

### Population-average test suggests non-exponential growth for *C. glutamicum*

A standard way of characterizing single-cell bacterial growth, is to determine the average relation between birth length *l*_b_ and division length *l*_d_ (Amir 2014). For *C. glutamicum*, we find an approximately linear relationship between these birth and division lengths, with a slope of 0.91±0.16 (2XSEM, Figure 3 A). This indicates that on a population level, *C. glutamicum* behaves close to the *adder* model, in which cells on average grow by adding a fixed length before dividing (Jun and Taheri-Araghi 2015; Amir 2014).

**Figure 3.**
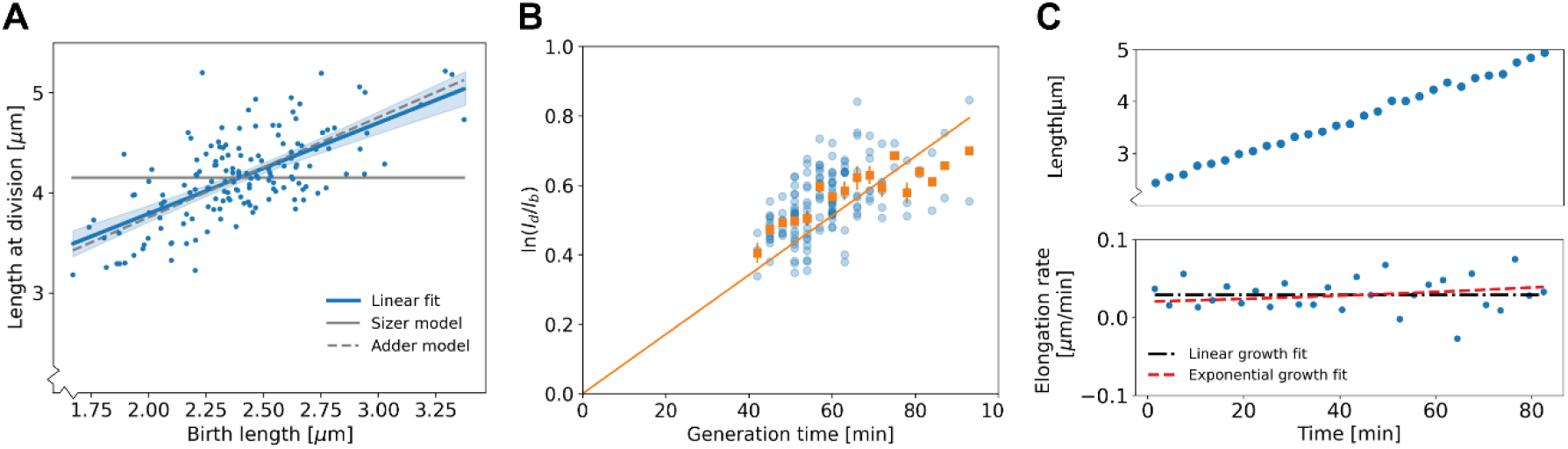
Population-level and single-cell level growth analysis. **(A)** Birth length *l*_b_ plotted against division length *l*_d_ for all measured cells, together with a linear fit (blue line), which has a slope of 0.91±0.16. Grey solid line: best fit assuming a pure sizer (slope 0). Grey dashed line: best fit assuming a pure adder (slope 1). The 95% confidence intervals of the linear fit, obtained via bootstrapping, are indicated by the blue shaded region. **(B)** Generation time versus 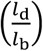 for all cells (blue dots) and the average per generation time (orange squares), with the standard error of the mean shown for all generation times for which at least 3 data points are available. The orange line represents a linear fit through the generation time averages that passes through the origin. For exponential growth, the averages would lie along this line, and the slope would be equal to the exponential growth rate. **(C)** Growth trajectory for a single cell (upper panel), together with its derivative for each measurement interval (lower panel). Fits to the derivative are shown for linear growth (black dash-dotted line) and exponential growth (red dashed line).

To investigate the growth dynamics from birth to division, we first tested if our cells conform to the generally observed exponential mode of single-cell growth. To this end, we applied a previously developed analysis on bacterial elongation data (Logsdon et al. 2017), by plotting 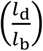 versus the growth time (Figure 3B). For an exponential grower, with the same exponential growth rate *α* for all cells, the averages of 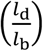 per growth time bin are expected to lie along a straight line with slope *α* intersecting the origin. By contrast, there appears to be a systematic deviation from this trend, with cells with shorter growth times lying above this line and cells with longer growth times lying below it, suggesting non-exponential elongation behavior. However, the significance and implications of these deviations for single-cell growth behavior are not clear from this analysis. There are several quantities that could be highly variable between cells that are averaged out in this representation, such as possible variations in exponential growth rate as a function of birth length, or variations in growth mode over time. Thus, a more detailed analysis of the growth trajectories is needed to rule out exponential growth, and to quantitatively characterize the growth dynamics.

The variability of key growth parameters is not easily extracted from individual growth trajectories due to the inherent stochasticity of the elongation dynamics and measurement noise (Figure 3C). In fact, it has been estimated that to distinguish between exponential and linear growth for an individual trajectory, the trajectory needs to be determined with an error of ~6% (Cooper 1998). Distinguishing subtler growth features may require an even higher degree of accuracy, which is presently experimentally unavailable (Appendix 3). Therefore, an analysis method is needed that is less noise-sensitive than an inspection of the single-cell trajectories, but simultaneously does not average out potentially relevant growth features such as time-dependence and birth length variability.

### Growth-inference method yields average elongation rate curves

To obtain quantitative elongation rate curves as a function of time and birth length, despite the high degree of individual variation, we developed a data analysis procedure that exploits the noise-reducing properties of multiple-cell conditional averaging. The key idea is to obtain an average dependence of the cellular length *L*(*t, l*_b_) on the time *t* since birth and birth length *l*_b_, by first obtaining the average dependence of *L*(*t, l*_b_) on *l*_b_ for each discrete value of *t* individually. This yields an average elongation curve for each birth length *l*_b_, without the need to perform inference on noisy *L*(*t*) single-cell curves.

The analysis procedure is as follows. First, for all cells in our data set, we determine the time since birth *t*, the cellular length *L* at time *t*, and the birth length *l*_b_. Subsequently, we relate the length at time *t* to the birth length, yielding a series of scatter plots for each measurement time (Figure 4A). Importantly, these scatterplots suggest a simple apparently linear relationship between *L* and *l*_b_. For each such plot, we thus make a linear fit through the data, yielding a family of curves *L_t_*(*l*_b_) for each time since birth *t* (Figure 4B). Higher-order fitting functions result in a negligible improvement of the goodness-of-fit, while increasing the mean error on inferred elongation rates (Appendix 2-Figure 3). Note that for both purely linear and purely exponential growth, *L*(*t, l*_b_) would depend linearly on *l*_b_: for linear growth *L*(*t, l*_b_) = *αt* + *l*_b_, whereas for exponential growth *L*(*t, l*_b_) = *l*_b_ exp(*αt*) (Appendix 2-Figure 6). From the family of relations *L_t_*(*l*_b_), we compute a series of points {*L*(*t*_0_, *l*_b_), *L*(*t*_1_, *l*_b_), *L*(*t*_2_, *l*_b_),…}, yielding the average growth trajectory of a cell starting out at length *l*_b_ (Figure 4C). Note, we must remove a bias in the *l*_b_ associated with each average trajectory, arising from measurement noise in the cell lengths at birth (Appendix 4). In summary, this procedure allows us to obtain an unbiased interference of the average elongation trajectories as a function of the cell’s birth length.

**Figure 4.**
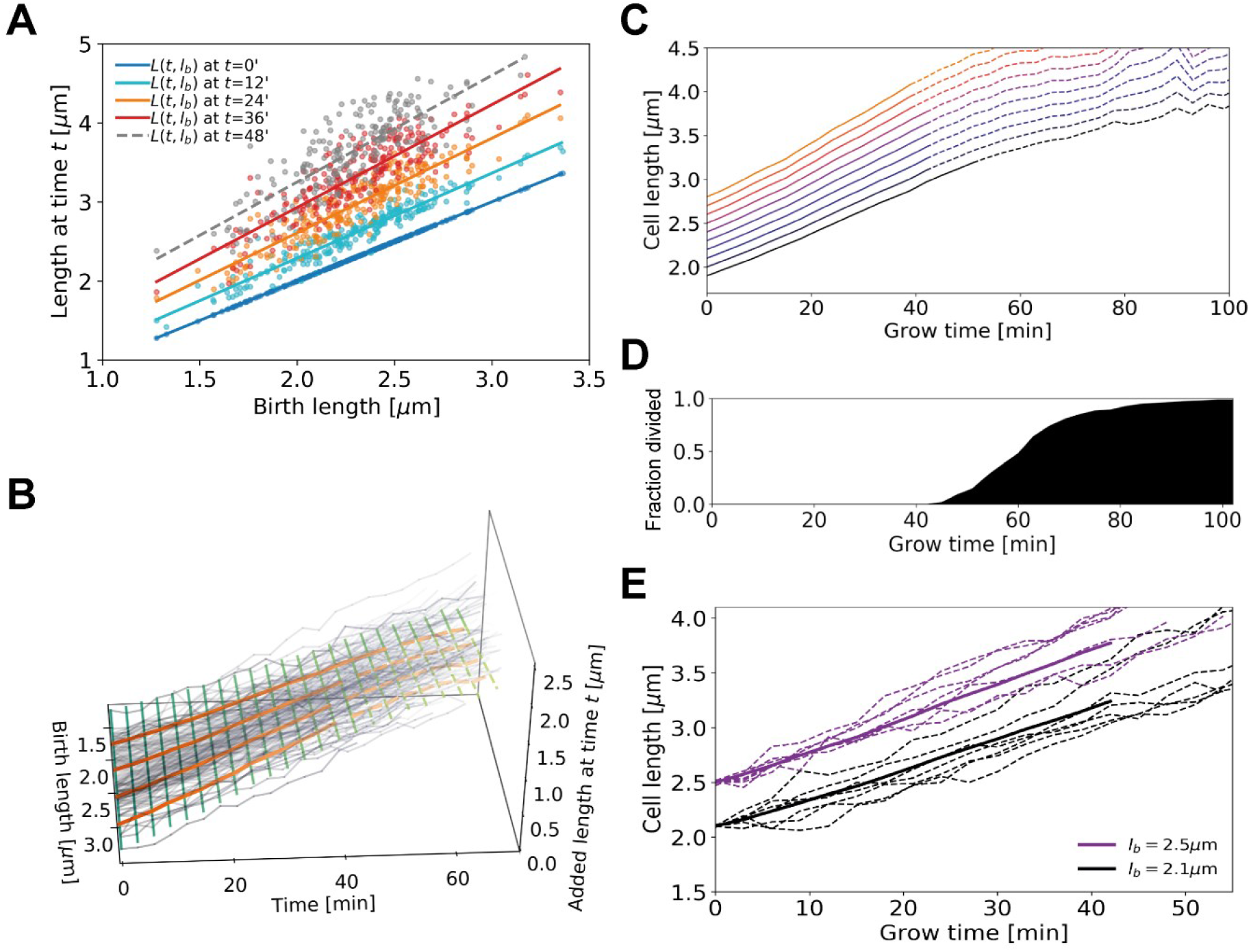
Average elongation curve inference procedure. **(A)** For each cell, the length *L*(*t*) at different times *t* since birth is plotted as a function of birth length *l_b_*. A linear fit of the resulting “wave front” is performed for each time *t*. This allows us to determine average cell length *L*(*t, l_b_*) at time *t* as a function of birth length *l_b_*. **(B)** 3D representation of the inference method of average length trajectories, with the added length *L*(*t, l_b_*) – *l_b_* on the z-axis. Elongation trajectories for individual cells are indicated in grey, linear fits through all cell lengths at each timestamp are indicated by green lines. The orange lines represent four sample average length trajectories, obtained by connecting all values of the green lines associated with one birth length. Dotted lines represent regimes where averages are biased due to dividing cells. **(C)** Average elongation trajectories obtained from the fits shown in (A) for a range of birth lengths, starting at 1.9 μm with steps of 0.1 μm (solid lines). The dashed lines represent regions where the inferred elongation curves are biased due to dividing cells, and are excluded from subsequent analysis. **(D)** Cumulative fraction of cells divided as a function of grow time. **(E)** Elongation trajectories for cells with birth lengths close to 2.5 μm (purple dashed lines) and birth lengths close to 2.1 μm (black dashed lines) together with their respective inferred average trajectories (purple solid line and black solid line).

### Elongation rate inference reveals asymptotically linear growth mode

Our inference approach yields the functional dependence of the average added length on growth time and birth length. We find that the average length steadily increases initially, but levels off and shows pronounced fluctuations for larger growth times (Figure 4C). This late-time behavior (dashed lines in Figure 4C) is caused by decreasing cell numbers due to division events (Figure 4D), which also introduces a bias in the averaging procedure. After the first division event, the average inferred growth would be conditioned on the cells that have not divided yet. For a given birth length, faster-growing cells divide earlier than slower-growing cells (Appendix 2-Figure 2) causing this conditional average to underestimate cellular elongation rates for the whole population after the first division. Because our aim is to infer elongation curves that characterize the whole population, ranging from slow to fast growers, for further analysis only the part of each trajectory before the first division event is used (Figure 4D). Sub-population elongation curves can also be obtained that extend past the first division event, but only if the entire analysis for these curves is performed only on these slower-dividing cells (Appendix 2-Figure 4).

We obtain elongation rate curves by taking a numerical derivative of smoothed growth trajectories (Appendix 5). To determine the associated error margins of the elongation rates, we use a custom bootstrapping algorithm (Efron 1979). The resulting 2σ bounds are shown as semitransparent bands. Despite the high noise level of individual elongation trajectories, the inferred average elongation rates have an error margin of around 8%. Thus, our approach robustly infers average elongation trajectories from single-cell growth data. Elongation rates of cells with larger birth length are consistently higher than the elongation rates of cells with smaller birth length. Strikingly, the elongation rate curves initially increase, but then gradually level off towards a linear growth mode (Figure 5). We note a slight difference in the cell elongation rates between the strain expressing DivIVA-mCherry (Figure 5A) and wild type cells (Figure 5B). Importantly, this difference does not qualitatively change the mode of growth, but does show that a translational fusion to DivIVA tends to lower elongation rates.

**Figure 5.**
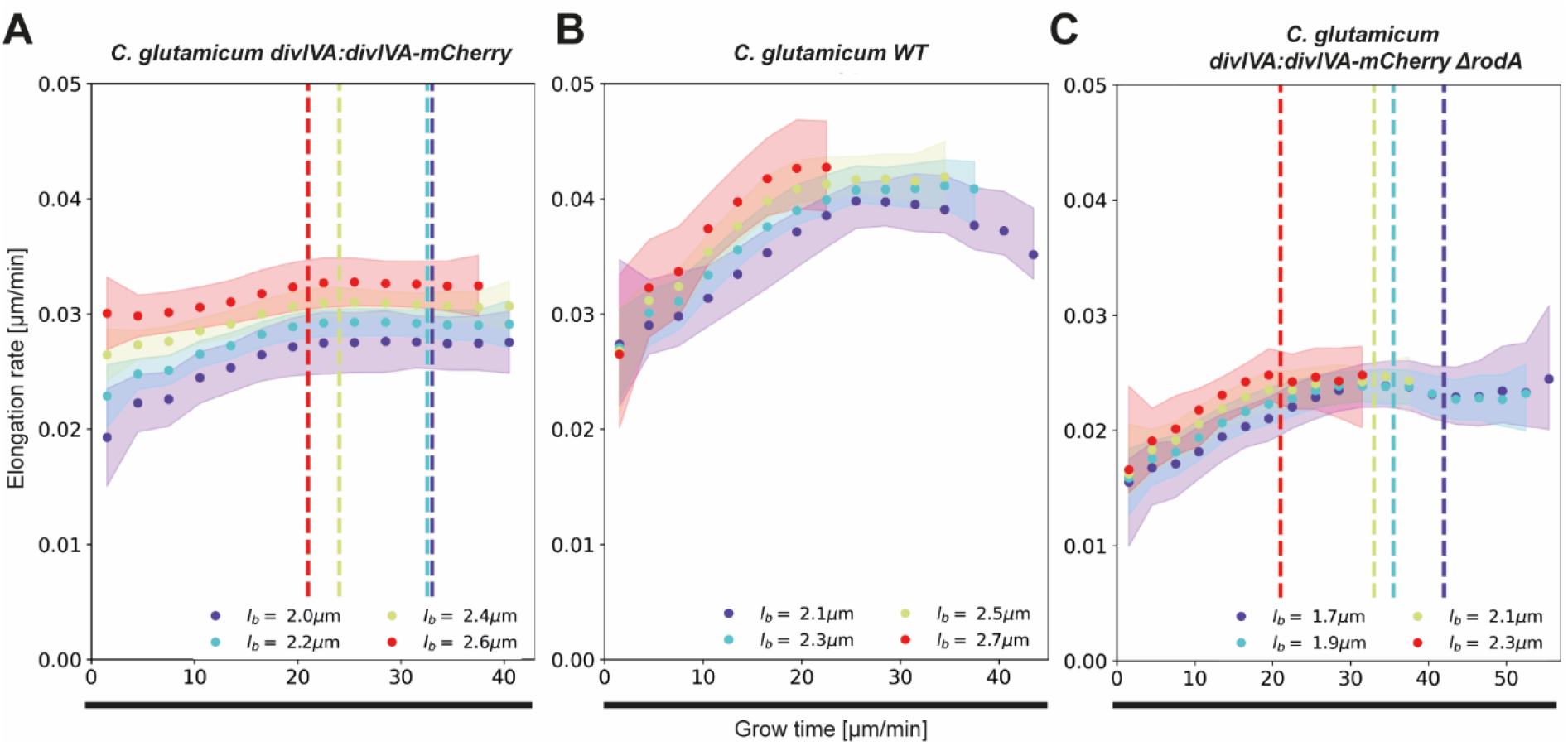
Inferred average elongation rates. (A) Average elongation rates for four birth lengths (dots), for the DivIVA labelled cells. The 2σ confidence intervals obtained by bootstrapping are indicated by the shaded areas. Vertical dashed lines: average onset of septum formation per birth length. **(B)** Average elongation rate trajectories for the wild-type cells, confidence intervals shown as in (A). **(C)** Average elongation rate trajectories for the *ΔrodA* mutant, confidence intervals shown as in (A).

To test the performance of our proposed inference method, we simulated a population of growing cells with a presumed growth mode from which we sample cells lengths as in our experiments, including measurement noise (Appendix 3). We ran simulations for cells performing linear growth, exponential growth, and the growth mode inferred here for DivIVA-labelled cells (Figure 5A). We find that our inference method is able to recover the input growth mode with high precision in all cases (Appendix 4, Appendix 6), demonstrating the accuracy and internal consistency of our inference method.

### Onset of the linear growth regime does not consistently coincide with septum formation

A central feature of the obtained elongation rate curves is a transition from an accelerating to a linear growth mode after approximately 20-25 minutes (Figure 5). One possibility is that this levelling off is connected with the onset of division septum formation. Given that the FtsZ-dependent divisome propagates the invagination of the septum under the consumption of cell wall precursors (e.g. Lipid-II), we hypothesized that the appearance of the additional sink for cell-wall building blocks could lead to coincidental leveling-off of the elongation rates (Scheffers and Tol 2015). To test this hypothesis, we used the moment of a sharp increase in the average DivIVA-mCherry signal at the cell center as a proxy for the moment of onset of septum formation (Appendix 2-Figure 7): the inward growing septum introduces a negative curvature of the plasma membrane, leading to the accumulation of DivIVA (Lenarcic et al. 2009; Strahl and Hamoen 2012). We observe that onset of septum formation does not consistently coincide with the moment at which the elongation rate levels off (Figure 5 A): for smaller cells the onset of septum formation occurs much later. Therefore, it seems implausible that the observed linear growth regime is due the septum acting as a sink for cell-wall building blocks.

### Polar cell wall formation is the rate-limiting step for growth, leading to a linear growth regime

To provide insight into the anomalous single-cell growth behavior, we model single-cell elongation as being rate-limited by the apical cell wall formation mechanism. To formulate this rate-limiting apical growth (RAG) model, we first consider the biochemical pathway that leads to cell wall formation in *C. glutamicum*, as illustrated in Figure 1. The key process for cell wall formation in *C. glutamicum* is polar peptidoglycan (PG) synthesis. PG intermediates are provided by the substrate Lipid-II, and the integration of new material into the PG-mesh is mediated by transglycosylases (TGs) located at the cell pole. At the TG sites, Lipid-II is translocated across the plasma membrane by the Lipid-II flippase MurJ (Sham et al. 2014; Kuk, Mashalidis, and Lee 2017; Butler et al. 2013). After PG building blocks provided by Lipid-II are incorporated into the existing cell wall by transglycolylation, transpeptidases (TP) conduct the crosslinking of peptide subunits, which contributes to the rigidity of the cell wall (Scheffers and Pinho 2005; Valbuena et al. 2007; Schleifer and Kandler 1972). During growth, the area of the PG sacculus, and thus the number of TG sites, is extended by RodA and bifunctional penicillin binding proteins (PBPs), recruited by DivIVA (Letek et al. 2008).

To model this growth mechanism, we assume that the rate of new cell wall formation is proportional to the number of TG sites *N*(*t*). We describe the interaction between Lipid-II and TG sites by Michaelis-Menten kinetics (Figure 6A). Specifically, if the cell length added per unit time is proportional to the cell wall area added per unit time, we find

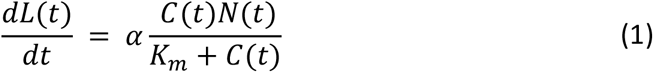

with *L*(*t*) the cell length at time *t, C*(*t*) the concentration of Lipid-II, *K_m_* the Michaelis constant for this reaction, and *α* is a proportionality constant.

**Figure 6.**
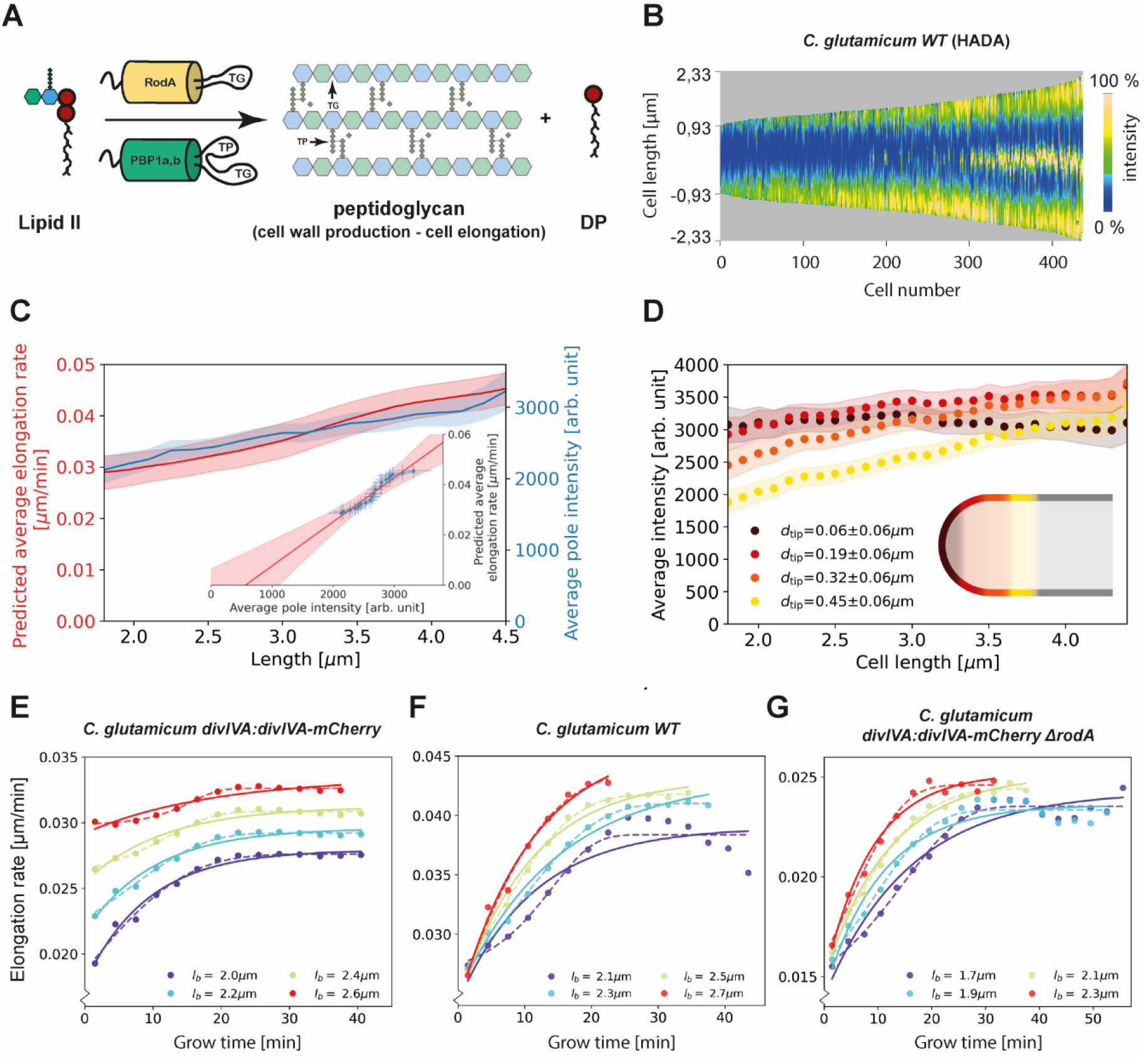
Modelling of average elongation rates using HADA staining results. **(A)** Schematic depicting cell wall formation via Lipid-II and transgrlycosylases (TG’s). The corresponding Michaelis-Menten equation describes the change of length over time as function of the Lipid-II concentration *C*(*t*) and the number of the TG sites *N*(*t*). **(B)** Demograph of *C. glutamicum* cells stained with HADA. Cell are ordered by length, with the stronger signal oriented downwards. **(C)** Average elongation rate as a function of cell length (red), predicted from obtained average elongation rate curves (Appendix 7), together with the average HADA staining intensity at the cell pole after background correction (blue). The cell pole is defined here as the region within 0.77 μm (60 pixels) of the cell tip. The shaded regions indicate the 2XSEM bounds. For both curves, a moving average over cells within 0.7 μm of each x-coordinate is applied over the underlying data. Inset: predicted average elongation rate versus average HADA staining intensity (blue dots). A linear fit through the result (red line) is consistent with a proportional relationship. **(D)** Average HADA intensity as a function of cell length, shown for four regions close to the cell tip. A moving average over cells within 0.7 μm of each x-coordinate is applied over the underlying data. **(E-G)** Dots: average elongation rate curves as shown in Figure 5A. Solid lines: best fit of elongation model from Eq. (2), which assumes constant transglycosylase recruitment. Dashed lines: best fit of elongation model from Eq. (3), which assumes an exponential increase of transglycosylase recruitment.

To gain insight into the cell-cycle-dependence of *N*(*t*) and *C*(*t*), we made use of the cyan fluorescent D-alanine analogue HADA (see Material and Methods) to stain newly inserted peptidoglycan. Exponentially growing *C. glutamicum* cells were labelled with HADA for 5 minutes before imaging. The HADA stain will mainly appear at sites of nascent PG synthesis. As expected, HADA staining resulted in a bright cyan fluorescent signal at the cell poles and at the site of septation. Still images were obtained with fluorescence microscopy and subjected to image analysis (Figure 2A, 6B, Material and Methods).

We first verify that the HADA intensity profile at the cell poles can be used as a measure for the peptidoglycan insertion rate. To do this, we assume that the HADA intensity profile has two relevant contributions: fluorescent probe present in the cell plasma, and fluorescent probe attached to newly inserted peptidoglycan. We use the minimum of the HADA intensity profile, consistently located around mid-cell, as an estimate of the contribution from the cell plasma in each cell, and subtract this from the entire cellular profile to obtain the corrected HADA profile (Appendix 2-Figure 8). We then define the polar regions where we use the corrected HADA intensity to measure newly inserted peptidoglycan as the portions of the cell within 0.78 μm of the cell tips. Our results are, however, not strongly dependent on this polar region definition (Appendix 2-Figure 10). Subsequently, we compute a moving average of the corrected polar HADA intensity as a function of cell length (Figure 6C). These polar HADA intensities are approximately proportional to the inferred average single-cell elongation rates (Appendix 7), as shown in the inset of Figure 6C. Since a proportional relationship between elongation rate and peptidoglycan insertion rate is expected, this supports our interpretation of the corrected HADA polar intensity as the peptidoglycan insertion rate.

Analyzing the HADA intensity profile for smaller segments within the polar region, we find that the increase in intensity is unevenly distributed (Figure 6D). Close to the cell tip, the HADA intensity remains approximately constant across cell lengths, whereas a linear increase over cell lengths is seen further from the tip. Considering the implications of these measured intensities for *C*(*t*) and *N*(*t*) within our model in Eq. (1), we argue for a scenario where either *C*(*t*) is constant or *C*(*t*) ≫ *K_m_*. Our reasoning is as follows. From Eq. (1), we see that the approximately constant intensity at the cell tip can be produced in two ways: **(1)** *C*(*t*) ≫ *K_m_* or *C*(*t*) is constant across cell lengths, and the number of transglycosylases at the tip *N*_tip_(*t*) is constant, or **(2)** *N*_tip_(*t*) and *C*(*t*) anticorrelate in such a way to produce constant insertion. However, we consider constant *N*_tip_(*t*) as biologically the most plausible scenario. This is supported by noting that the concentration of Lipid-II is the same directly before and after division, such that *C*(*t*), and by implication *N*_tip_(*t*), is similar for the shortest and the longest cell lengths (Appendix 2-Figure 9). In our subsequent analysis, we will therefore assume that either *C*(*t*) is constant, or *C*(*t*) ≫ *K_m_*. This implies that 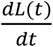 in Eq. (1) is directly proportional to *N*(*t*).

To derive an expression for *N*(*t*), we first note that the old and new cell pole in the cell need to be treated differently. We assume the number of polar TG-sites to saturate within one cellular lifecycle, such that the new pole initiates with *N*(*t*) below saturation, while the old pole - inherited from the mother cell - is saturated. Letting the number of TG sites increase proportional to the number of available sites, we arrive at the following kinetic description for *N*(*t*):

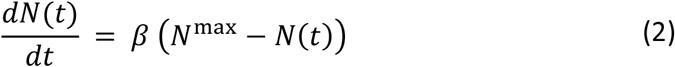

Here, *N*_max_ is the maximum number of sites at the cell poles, and *β* is a rate constant. This result, together with Eq. (1), defines our RAG model. The predicted elongation rates provide a good fit to the experiment for all studied genotypes (Figure 6E-G), although the data appear to exhibit a stronger inflection.

Instead of assuming a constant recruitment of TG enzymes, we can construct a more refined model that takes TG recruitment dynamics into account. There is evidence that transglycosylase RodA and PBPs are recruited to the cell pole via the curvature-sensing protein DivIVA (Letek et al. 2008; Sieger et al. 2013). As shown in (Lenarcic et al. 2009), DivIVA also recruits itself, leading to the exponential growth of a nucleating DivIVA cluster. Therefore, we let the recruitment rate of TG enzymes be proportional to the number of DivIVA proteins *N*_D_(*t*) = *N*_D_(0)*e^γt^*. This results in a modified kinetic description for *N*(*t*) (Eq. (2)):

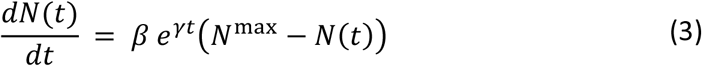

This refined model can capture more detailed features of the measured elongation rate curves (Figure 6E-G), including the stronger inflection, with an additional free parameter, *γ*, encoding the self-recruitment rate of DivIVA.

The central assumption of our RAG model is that the growth of the cell poles, mediated via accumulation of TG enzymes, is the rate-limiting step for cellular growth. To test this assumption, we repeated our experiment with a *rodA* knockout (Sieger et al. 2013). The SEDS-protein RodA is a mono-functional TG (Meeske et al. 2016; Emami et al. 2017; Sjodt et al. 2018), whose deletion results in a phenotype with a decreased population growth rate in the shaking-flask (Sieger et al. 2013). The cells’ viability is nonetheless backed up by the presence of bifuncional class A PBPs capable of catalyzing transglycoslyation and transpeptidation reactions. We expect this knockout to lower the efficiency of polar cell wall formation, thus slowing down the rate-limiting step of growth. Specifically, we expect the knockout of *rodA* to mainly affect the efficiency of Lipid-II integration into the murein sacculus. Within our RAG model, this translates to a lowering of the cell wall production per transglycosylase site *α*. This would imply elongation rate curves of similar shape for the *ΔrodA* mutant, only scaled down by a factor *α*^WT^/*α*^ΔrodA^. Indeed, we observe such a scaling down of the elongation rate curves (Figure 5C), lending further credence to our model for *C. glutamicum* growth.

A striking feature observed across growth conditions and birth lengths, is the onset of a linear growth regime after approximately 20 minutes (Figure 5A-C). The robustness of this timing can be understood from the RAG model: the regime of linear growth is reached via an exponential decay of the number of available TG sites until saturation is reached. This exponential decay makes the moment of onset of the linear growth regime relatively insensitive to variations in *N*(0) and *N*^max^. Specifically, from Eq. (2), it can be shown that the difference between *N*(*t*) and *N*^max^ is halved every ln(2)*β* minutes, which amounts to ~8 minutes given fitted value of *β* (Appendix 8-Table 1).

Finally, our RAG model makes a prediction for the degree of transglycosylase saturation of the cell poles at birth, relative to the saturation in the linear growth regime. We find that this saturation is comparable between wild-type and the *ΔrodA* mutant (~65% on average), but significantly higher for DivIVA labelled cells (~80% on average) (Appendix 8 Tables 1 and 2). This suggests that the number of transglycosylase sites at birth is relatively high in the DivIVA labelled cells.

### Birth length distribution of linear growers is more robust to single-cell growth variability

After obtaining average single-cell growth trajectories, we next asked how this growth behavior at the single cell level affects the growth of the colony. It was shown that asymmetric division and noise in individual growth times results in a dramatic widening of the cell-size distribution for a purely exponential grower (Marantan and Amir 2016). For an asymptotically linear grower, however, we would expect single-cell variations to have a much weaker impact.

To quantify the difference between asymptotically linear growth and hypothetical exponential growth for *C. glutamicum*, we performed population growth simulations for both cases. For the asymptotically linear growth, we assumed the elongation rate curves obtained from our model. For exponential growth, we assumed the final cell size to be given by *l*_d_ = *l*_b_ exp(*α*(*t_t_* + Δ*t*)) + Δ*l*, with *α* the exponential elongation rate, *t_t_* the target growth time, Δ*t* a time-additive noise term and Δ*l* a size-additive noise term. All growth parameters necessary for the simulation were obtained directly from the experimental data (Appendix 9). From this simulation, the distribution of initial cell lengths was determined for each scenario.

The resulting distribution of birth lengths for the asymptotically linear growth case closely matches the experimentally determined distribution (Figure 7). By contrast, the distribution for exponential growth is much wider, and exhibits a broad tail for longer cell lengths. This suggests a strong connection between growth mode and the effect of individual growth variations on population statistics. *C. glutamicum* has a high degree of variation of division symmetry (Appendix 9-Figure 1C) and single-cell growth times, but due to the asymptotically linear growth mode, the population-level variations in cell size are still relatively small.

**Figure 7.**
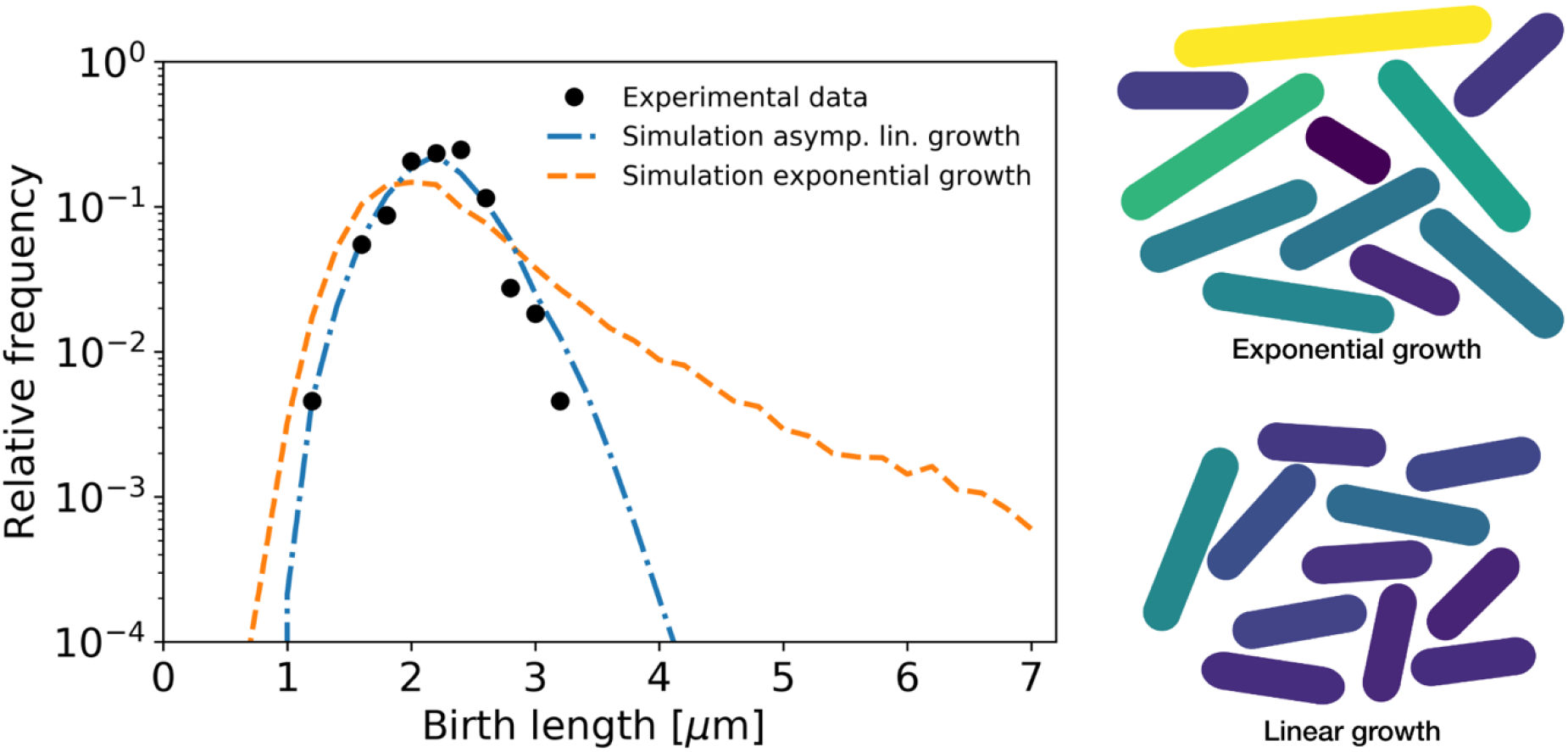
Simulation of population growth for asymptotically linear and exponential growth. Left: birth length distribution for simulated asymptotically linear growth (blue dash-dotted line), and for simulated exponential growth (orange dashed line). For both simulations, all relevant growth parameters and distributions are obtained directly from the experimental data. Black dots: experimental birth length distribution. Right: sample of 11 cells from the exponential and asymptotically linear growth simulations, color coded according to length.

## Discussion

By developing a novel growth trajectory inference and analysis method, we showed that *C. glutamicum* exhibits asymptotically linear growth, rather than the exponential growth generally assumed for most species. The obtained elongation rate curves are shown to be consistent with a model of apical cell wall formation being the rate-limiting step for growth. The RAG model is further validated by experiments with a *ΔrodA* mutant, in which the elongation rate curves look functionally similar, but with a downward shift compared to wild type (Figure 5B, C), as expected based on our model. For *C. glutamicum*, apical cell wall formation is a plausible candidate for the rate-limiting step of growth, because synthesis of the highly complex cell wall and lipids for the mycolic acid membrane is cost intensive and a major sink for energy and carbon in *Corynebacteria* and *Mycobacteria* (Brennan 2003). An analysis of elongation rates as a function of time and birth length has previously been done in *B. subtilis* by binning cells based on birth length (Nordholt, van Heerden, and Bruggeman 2020). Applying this method to our data set yields elongation rates averaged over cells within a binning interval (Appendix 2-Figure 5). Averaging our inferred elongation rates over the same bins, we find the two methods to yield consistent results. The binning method however involves a tradeoff: a smaller bin width results in a larger error on the inferred elongation rates, whereas a larger bin width averages out all variation within a larger birth length interval. Our method does not suffer from this binning-related tradeoff, and it provides detailed elongation rate curves at any given birth length.

Our proposed growth model shares some similar features to recent experimental observations on polar growth in Mycobacteria (Hannebelle et al. 2020). Polar growth was shown to follow ‘new end take off’ (NETO) dynamics (Hannebelle et al. 2020), in which the new cell pole makes a sudden transition from slow to fast growth, leading to a bilinear polar growth mode. In our proposed growth model for *C. glutamicum* however, the new pole gradually increases its average elongation rate before saturating to a constant maximum. The deviation of *C. glutamicum* from NETO dynamics can also be seen by comparing each of the pole intensities in the HADA staining experiment, which does not show any signatures of NETO-like growth (Appendix 2-Figure 11). It remains unclear which molecular mechanisms produce the differences in growth between such closely related species. However, the mode of growth described here for *C. glutamicum* might well be an adaption to enable higher growth rates.

To investigate the implications of our inferred single-cell growth mode for cell-size homeostasis throughout a population of cells, we performed simulations of cellular growth and division over many generations. We found that our asymptotically linear growth model accurately reproduces the experimental distribution of cell birth lengths. By contrast, a model of exponential growth predicts a much broader distribution with a long tail for larger birth lengths. This indicates a possible connection between mode of growth and permissible growth-related noise levels for the cell. Indeed, if single-cell growth variability is reduced by a factor 3, the distributions corresponding to both growth modes show a similarly narrow width (Appendix 9-Figure 2). However, an asymptotically linear grower is able to maintain a narrow distribution of cell sizes even for higher noise levels, whereas for an exponential grower this distribution widens dramatically (Figure 7).

The enhanced robustness of the length distribution of linear growers is interesting from an evolutionary point of view. Most rod shaped bacteria use sophisticated systems, such as the Min system, to ensure cytokinesis precisely at midcell (Bramkamp and van Baarle 2009; Lutkenhaus 2007). Bacteria encoding a Min system grow by lateral cell wall insertion. In contrast, rod-shaped bacteria in the *Actinobacteria* phylum such as *Mycobacterium* or *Corynebacterium* species, grow apically and do not contain a Min system, nor any other known division site selection system (Donovan and Bramkamp 2014). *C. glutamicum* rather couples division site selection to nucleoid positioning after chromosome segregation via the ParAB partitioning system (Donovan et al. 2013), and has a broader distribution of division symmetries. We speculate that due to *C. glutamicum’s* distinct growth mechanism, a more precise division site selection mechanism is not necessary to maintain a narrow cell size distribution.

The elongation rates reported in this work reflect the increase in cellular volume over time. However, the increase in cell *mass* is not necessarily proportional to cellular volume. In exponentially growing *E. coli*, the cellular density was recently reported to systematically vary during the cell cycle, while the surface-to-mass ratio was reported to remain constant (Oldewurtel et al. 2019). It is unknown how single-cell mass increases in *C. glutamicum*, but it would follow exponential growth if mass production is proportional to protein content. This raises the question how linear volume growth and exponential mass growth are coordinated. The presence of a regulatory mechanism for cell mass production that couples to cell volume is implied by the elongation rate curves obtained for the *ΔrodA* mutant. As the elongation rate is lower in this mutant, average mass production needs to be lowered compared to the WT in order to prevent the cellular density from increasing indefinitely.

Our growth trajectory inference method is not cell-type specific, and can be used to obtain detailed growth dynamics in a wide range of organisms. The inferred asymptotically linear growth of *C. glutamicum* is a stark deviation from the generally observed exponential single-cell bacterial growth, and suggests the presence of novel growth regulatory mechanisms.

## Experimental Procedures

### Culture and live-cell time-lapse imaging

Exponentially growing cells of *C. glutamicum divIVA::divIVA-mCherry* and *C. glutamicum divIVA::divIVA-mCherry ΔrodA* respectively, grown in BHI–medium (Oxoid) at 30°C and 200 rpm shaking, were diluted to an OD_600_ of 0.01. According to the manufacturer’s manual cells were loaded into a CellASIC-microfluidic plate type B04A (Merck Milipore) and mounted on a Delta Vision Elite microscope (GE Healthcare, Applied Precision) with a standard four-color InSightSSI module and an environmental chamber heated to 30°C. Images were taken in a three-minute interval for 10 h with a 100×/1.4 oil PSF U-Plan S-Apo objective and a DS-red-specific filter set (32% transmission, 0.025 s exposure).

### Staining of newly inserted peptidoglycan and visualization in demographs

For the staining of nascent PG, 1 ml of exponentially growing *C. glutamicum ATCC 13032* cells, cultivated in BHI–medium (Oxoid) at 30°C and 200 rpm, were harvested, washed with PBS and resuspended in 25 μl PBS, together with 0.25 μl of 5 mM HADA dissolved in DMSO. The cells were incubated at 30 °C in the dark for 5 minutes, followed by a two-time washing step with 1 ml PBS and finally resuspended in 100 μl PBS. To obtain still-phase-contrast and fluorescent micrographs, 2 μl of the cell suspension were immobilized on an agarose pad. For microscopy, an Axio Imager (Zeiss) equipped with EC Plan-Neofluar 100x/1.3 Oil Ph3 objective and a Axiocam camera (Zeiss) was used together with the appropriate filter sets (ex: 405 nm; em: 450 nm). For single-cell analysis and the visualization in demographs, custom algorithms, developed in FIJI and R (Schindelin et al. 2012)(R Development Core Team 2003), were used. The code is available upon request.

### Image analysis

For image analysis a custom made algorithm was developed using the open-source programs FIJI and R (Schindelin et al. 2012)(R Development Core Team 2003). During the workflow unique identifiers to single cell cycles are assigned. The cell outlines are determined manually. Individual cells per timeframe are extracted then from the raw image and further processed automatically. The parameters length, area and relative septum position are extracted and stored together with the genealogic information and the timepoint within the respective cell cycle. The combination of image analysis and cell cycle dependent data structuring yields a list that serves as a base for further analysis. The documented code is available at: https://github.com/Morpholyzer/MorpholyzerGenerationTracker

## ACKNOWLEDGMENTS

This work was further funded by grants from the Deutsche Forschungsgemeinschaft (project P05in TRR174, granted to M.B. and project P06 in TRR174, granted to C.B.). J.M. is supported by a DFG fellowship within the Graduate School of Quantitative Biosciences Munich (QBM). We thank our colleagues from C.B. and M.B. groups for discussions, feedback and comments on the manuscripts.

## Author contribution

J.M., F.M., M.B. and C.B. conceived the project and designed the experiments. J.M., F.M. performed theoretical and experimental parts, respectively. J.M., F.M., M.B. and C.B. analyzed the data and models and wrote the manuscript.

