## Supplemental Material for "Single-cell growth inference of *Corynebacterium glutamicum* reveals asymptotically linear growth"

[*] These authors contributed equally

+ Corresponding authors:

Chase Broedersz

### Appendix 1

**Single-cell growth mode for apical cell wall formation as a rate-limiting step for growth**

To study growth limited by polar cell wall formation, we start by considering the Michaelis-Menten equation describing this formation process (Main Text Eq. (1)):

$$\frac{dL(t)}{dt} = \alpha \frac{C(t)N(t)}{K_m + C(t)}, \tag{A1}$$

with $L(t)$ the cell length at time $t$, $C(t)$ the concentration of cell wall building blocks in the cytosol, $N(t)$ the number of transglycosylases at the cell pole, $K_m$ the Michaelis constant for this reaction, and α a proportionality constant.

In Main Text Figure 1, we consider two scenarios. **(1)** Abundant availability of cell wall building blocks, i.e. $C(t) \gg K_m$, and **(2)** scarcity of cell wall building blocks, i.e. $C(t) < K_m$.

**A1.1 Building block insertion as a rate-limiting step for growth**

In scenario (1), Eq. (A1) reduces to $\frac{dL(t)}{dt} = \alpha N(t)$. In the regime of a constant number of transglycosylases at the pole, this implies that $\frac{dL(t)}{dt}$ is constant, resulting in linear growth.

**A1.2 Building block availability as a rate-limiting step for growth**

In scenario (2), the dynamics of building block creation, usage, and dilution need to be considered to determine the cellular elongation rate behavior. For the number of building blocks in the cytosol as a function of time $n(t)$, we can write the following differential equation:

$$\frac{dn(t)}{dt} = aV(t) - b\frac{dV(t)}{dt}. \tag{A2}$$

Here, $a$ encodes building block production rate per unit volume, and $b$ encodes building block usage by the cell wall formation mechanism, making use of $\frac{dA(t)}{dt} \propto \frac{dV(t)}{dt}$. To connect Eq. (A2) to Eq. (A1), we note that $C(t) = \frac{n(t)}{V(t)}$. Restricting ourselves to the regime $C(t) \ll K_m$, we can rewrite Eq. (A1) to

$$\frac{dV(t)}{dt} = c\frac{n(t)}{V(t)}, \qquad \text{(A3)}$$

where we made use of $\frac{dL(t)}{dt} \propto \frac{dV(t)}{dt}$. Here, $c$ encodes the proportionality between volume increase and the concentration of building blocks.

Combining Eq. (A2) with Eq. (A3), we obtain a set of coupled nonlinear differential equations governing the time-evolution of $V(t)$. These equations have no simple analytic solution; however, we can numerically explore the dependence of $V(t)$ on the differential equation parameters. To do this, we first absorb $c$ into $n(t)$, leaving us with two free parameters and two boundary conditions. The boundary conditions we set by imposing $V(0) = 1$ and $V(1) = 2$. In Appendix 1-Figure 1A, we see that depending on the choice for $a$ and $b$ we can have either sublinear, approximately linear, or superlinear growth. This demonstrates that the single-cell growth mode is dependent on the physiology of building block creation and depletion in the cell.

We can further constrain the solution space by demanding that the concentration of building blocks $C(t) = \frac{n(t)}{V(t)}$ is the same at birth and division. In this scenario, the observed variation in elongation curves is smaller (solid lines Appendix 1-Figure 1B), however the corresponding elongation rates (dashed lines Appendix 1-Figure 1B) still show marked qualitative differences between parameter choices.

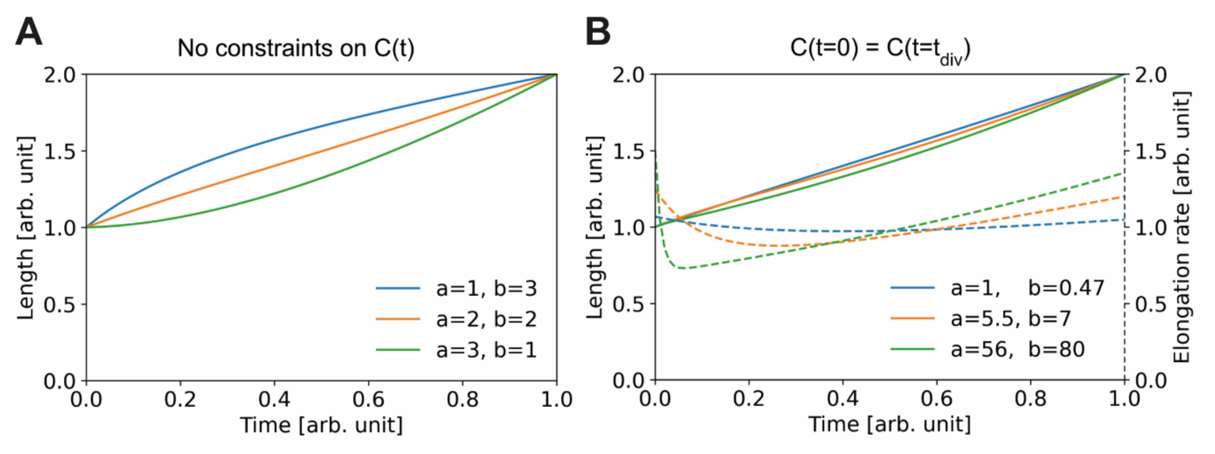


**Appendix 1-Figure 1 Elongation curves assuming building block availability is the limiting step for growth. (A)** Numerically obtained solutions for $V(t)$, from the set of coupled differential equations Eq. (A2) and Eq. (A3). For all solutions, $V(0) = 1$ and $V(1) = 2$ are imposed. **(B)** Solutions as in (A), but with the additional constraint that the concentrations before and after division are the same, i.e. $C(t = 0) = C(t = t_{\text{div}})$. Solid lines: solutions for $V(t)$. Dashed lines: corresponding $\frac{dV(t)}{dt}$, which are proportional to the concentration $C(t)$ per Eq. (A3).

#### Appendix 2: Supplementary figures

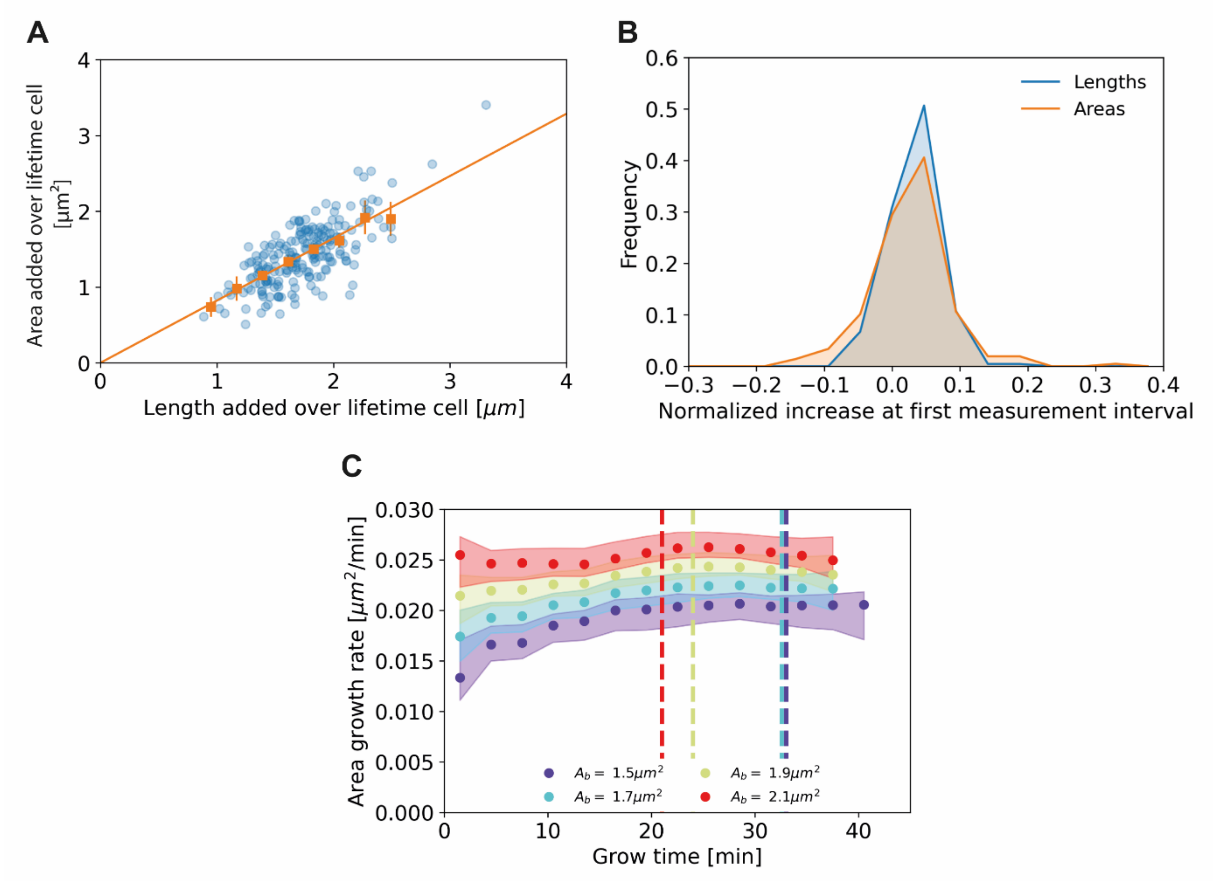


**Appendix 2-Figure 1 (A)** Length added versus area added over the cell lifetime for all cells included in our analysis (blue dots), together with averaged values at 0.2 µm intervals (orange squares) and 95% confidence intervals (orange vertical lines). The results are consistent with a proportional relationship (orange line). **(B)** Histogram of the normalized increase at first measurement interval using cell lengths (blue) and areas (orange). For the cellular lengths, this quantity is defined as $\frac{L(t=3min)-L(t=0)}{<l_b>}$, whereas for the areas it is defined as $\frac{A(t=3min)-A(t=0)}{<A_b>}$, with $A(t)$ the area at time $t$ and $A_b$ the birth area. The wider distribution for the areas suggests a higher measurement noise for this quantity. **(C)** Area growth rate for DivIVA labelled cells using estimated cell areas**.** The trajectories are consistent with those obtained from cell lengths (Main Text Figure 5A).

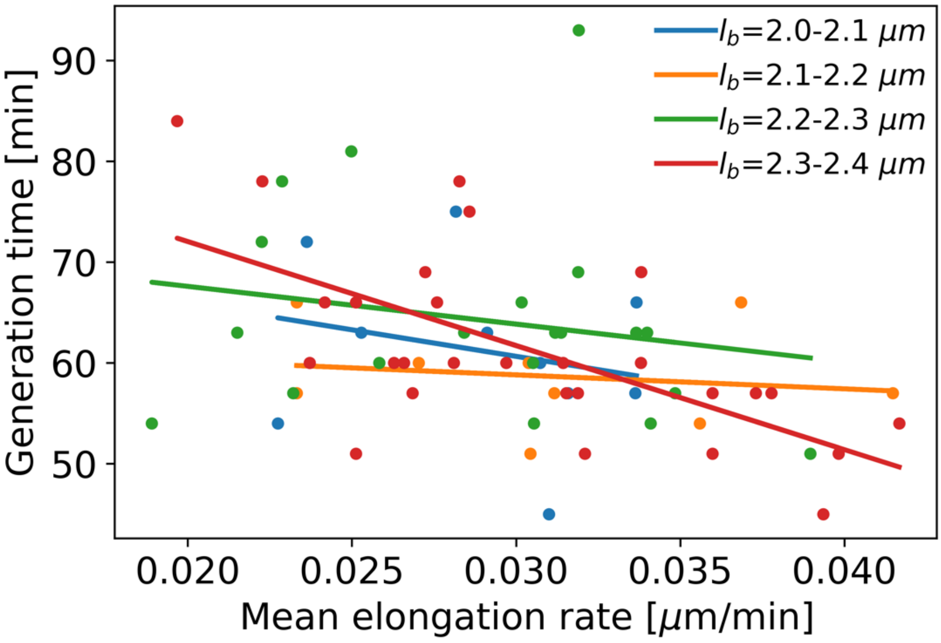


**Appendix 2-Figure 2** Mean elongation rate versus generation time for cells in four different birth size bins. Linear fits are indicated by solid lines. As generation times within a birth size bin tend to be shorter for faster-growing cells, the elongation rate curves obtained with our method become biased after the first division event. This justifies only using the part of the elongation rate curves until the first division event for further analysis.

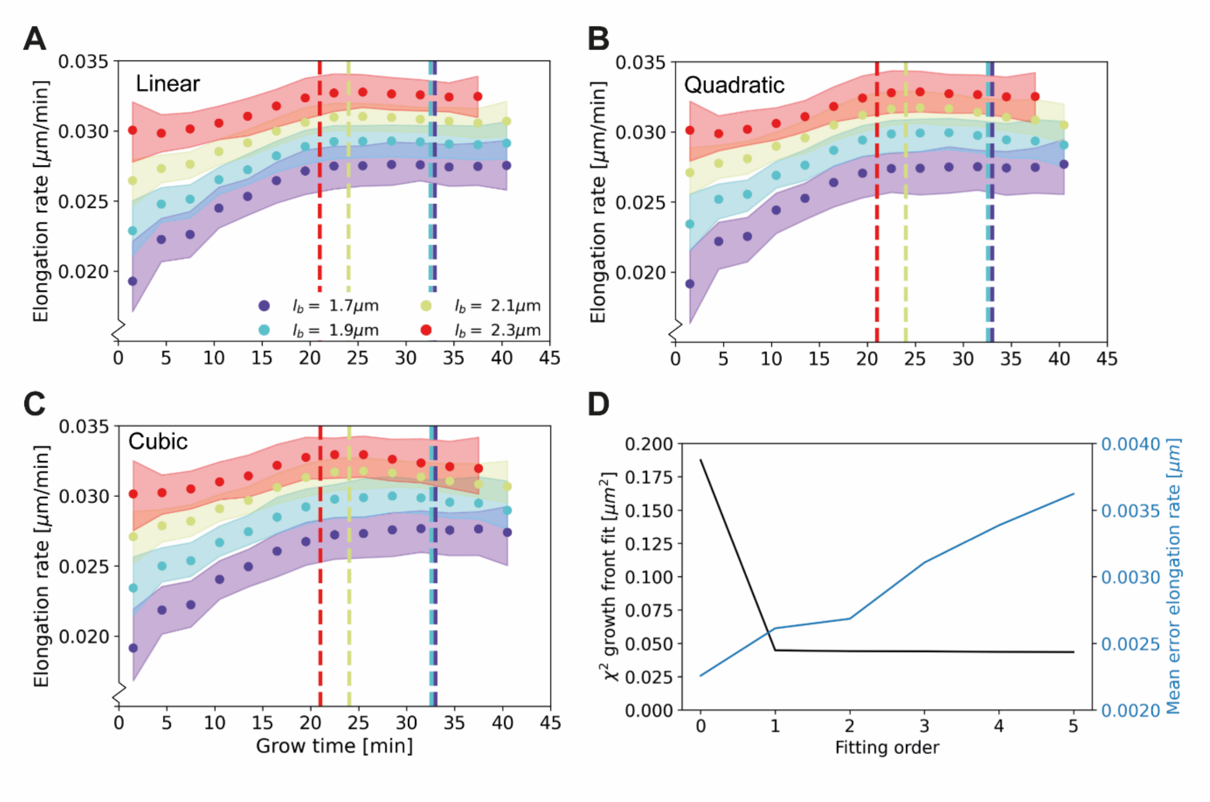


**Appendix 2-Figure 3** Elongation rate curves for different orders of the wave front fit of Main Text Figure 3A: Linear **(A)**, quadratic **(B)**, and cubic **(C)**. **(D)** $\chi^2$ of the fit of the wave front of Main Text Figure 3A for different fitting orders, together with the mean error on the elongation rate curves. The negligible improvement of the goodness-of-fit after the first order justifies the use of a linear fit for further analysis.

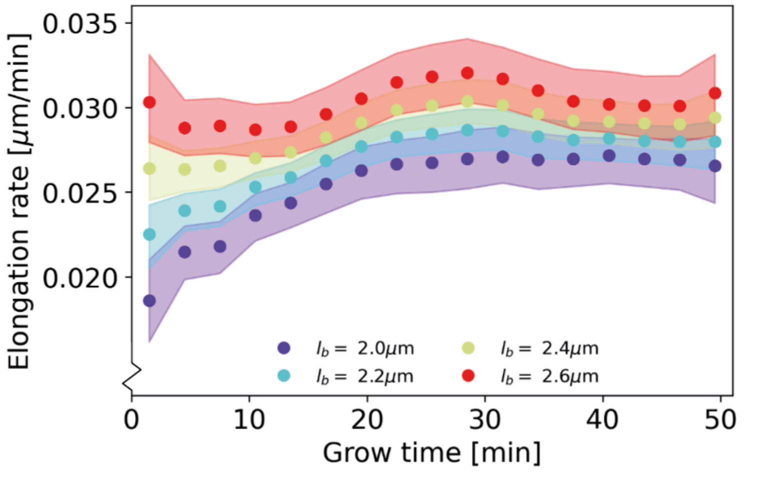


**Appendix 2-Figure 4** Conditional elongation rate curves**,** conditioned on DivVIA labelled cells that have a generation time of at least 51 minutes. The inferred elongation rate curves still display similar growth behavior to the unconditioned population, but exhibit a downwards shift, (Main Text Figure 5A). In addition, the linear growth phase observed until the cutoff time for the unconditioned population persists for longer grow times. Number of cells included: 139.

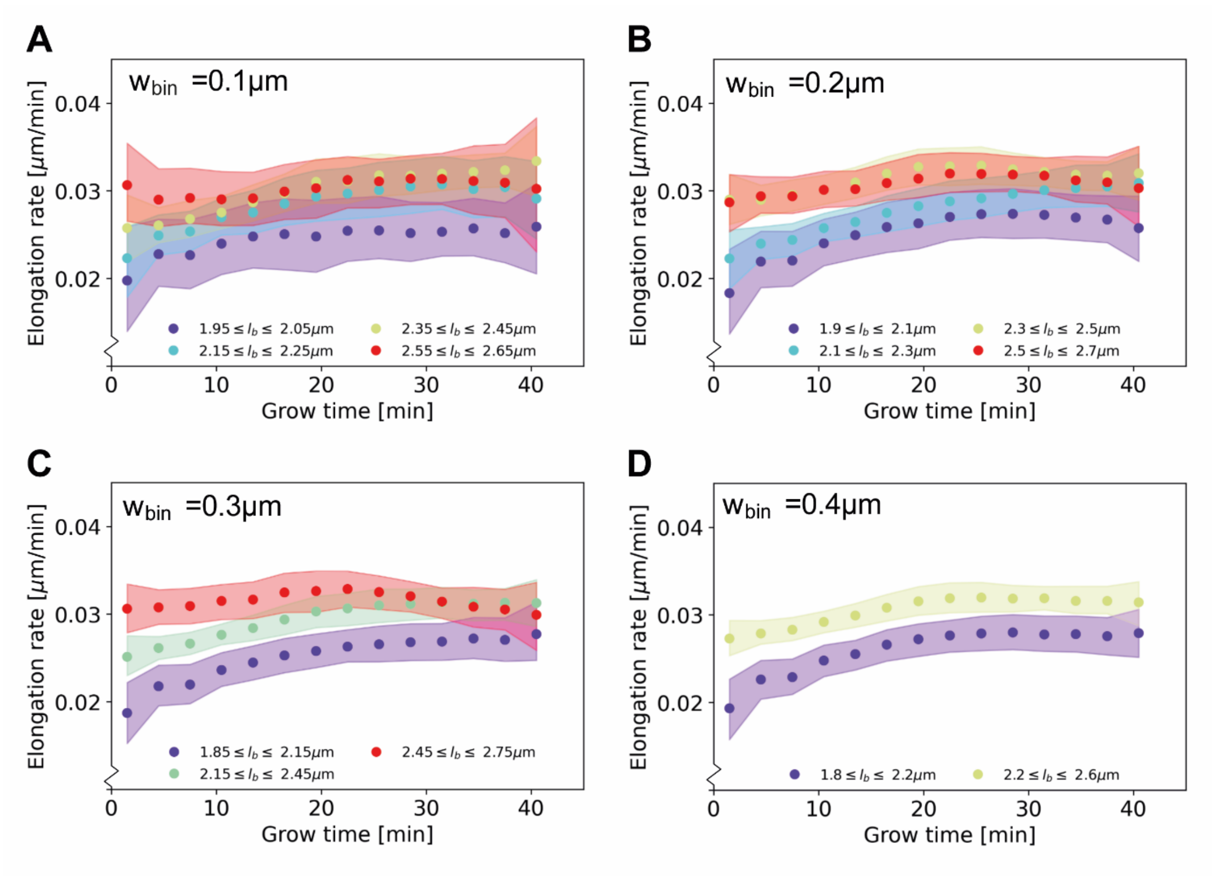


**Appendix 2-Figure 5** Elongation rate curves obtained through a binning procedure. Cells are divided into birth length bins, and for each bin the average length as a function of grow time is calculated. The resulting elongation curves are smoothened according to the same procedure as the elongation curves presented in the main text (see Appendix 5). From the smoothened elongation curves, elongation rates are calculated as a function of grow time. Results are shown for a bin width of 0.1 μm (**A**), 0.2 μm (**B**), 0.3 μm (**C**), 0.4 μm (**D**), where each $l_b$ indicates the center of the birth length bin.

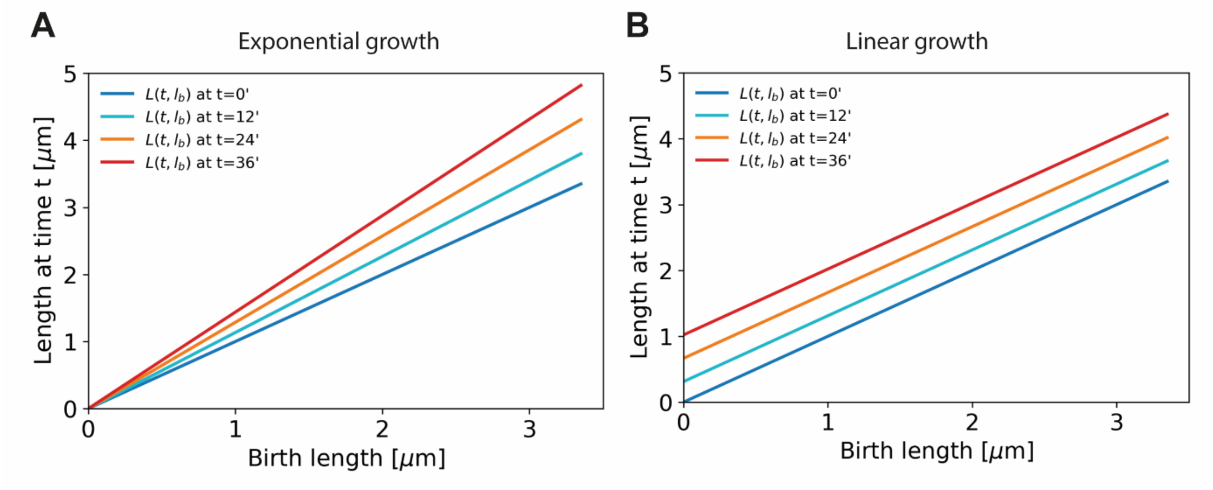


**Appendix 2-Figure 6** A linear fit through the cell lengths at each time step would be enough to describe exponential growth (**A,** offset is zero for all time stamps) as well as linear growth (**B,** slope is equal to 1 for all time stamps).

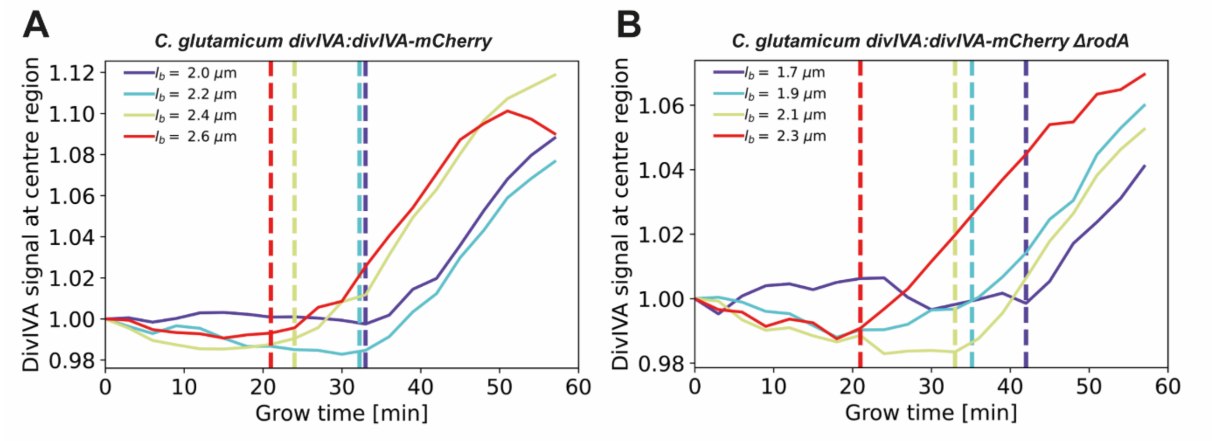


**Appendix 2-Figure 7** The average DivIVA-mCherry signal from the cell center over time is shown for DivIVA labelled cells (**A**) and ΔrodA DivIVA labelled cells (**B**). The cell center is here defined as the region between 20% and 80% of the total cell length. The onsets of septum formation, derived from the DivIVA signal-mCherry signal, are indicated by the dashed lines; these do not consistently coincide with the levelling off of elongation rates (Main Text Figure 5A). This is inconsistent with the leveling off being due to a competition between polar growth and septum formation.

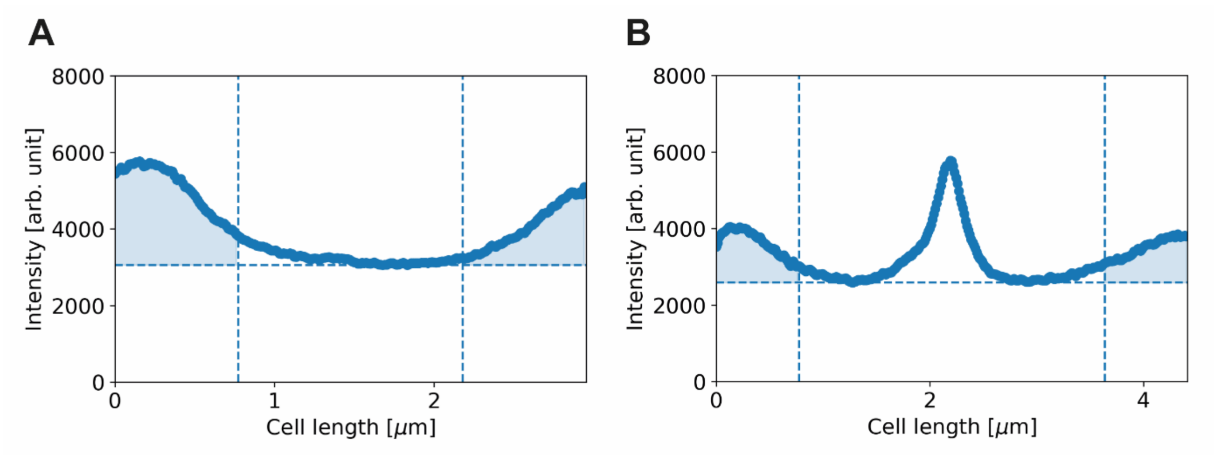


**Appendix 2-Figure 8 Calculation of corrected polar HADA intensity, illustrated for two HADA profiles.** Solid line: HADA intensity profile. Dashed horizontal line: minimum of HADA profile. Dashed vertical lines: boundary of polar region. Shaded area: calculated total polar intensity. Results shown for a cell with a length of 2.3 μm (**A**) and 4.4 μm (**B**).

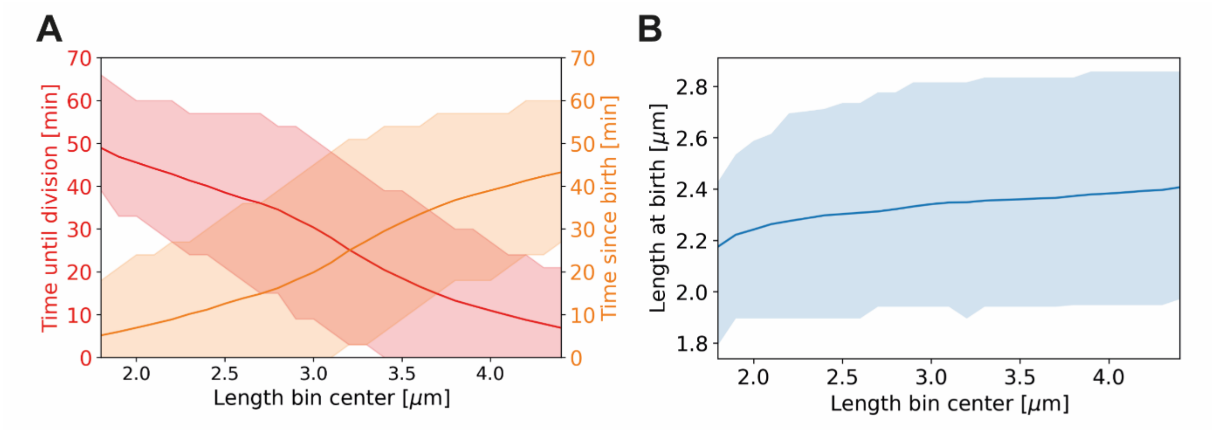


**Appendix 2-Figure 9 Average properties of wild-type cells as a function of length.** Values are shown over the range of observed lengths in the HADA staining experiment, using a moving average with the same width (±0.7 $\mu$m) as in Main Text Figure 6C. **(A)** Red line: average time until division, together with the two standard deviation bounds (red shaded area). Orange line: average time since birth, together with two standard deviation bounds (orange shaded area). **(B)** Blue line: average birth length for each birth length bin (blue line), together with the two standard deviation bounds (blue shaded area).

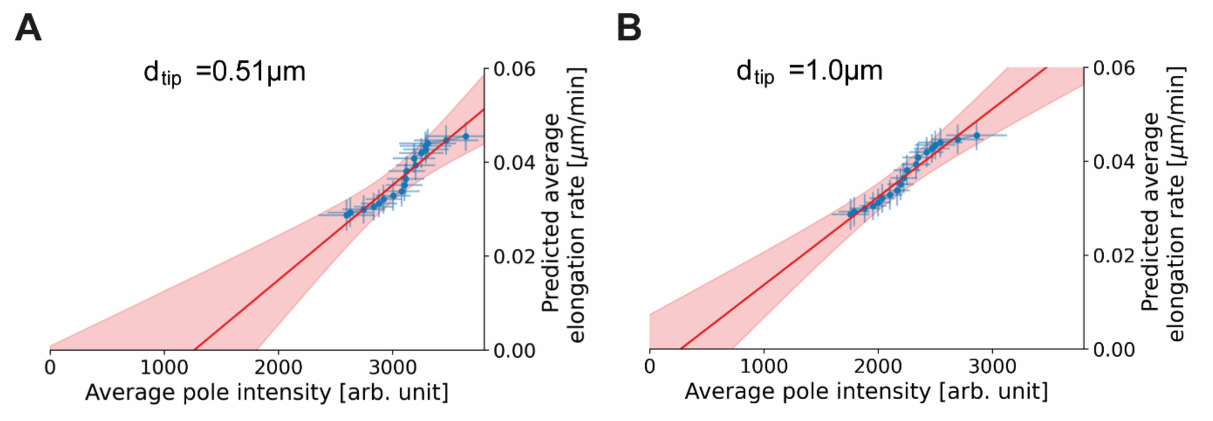


**Appendix 2-Figure 10 Proportionality between average pole intensity and predicted average elongation rate for different polar region definitions.** Average elongation rate as a function of cell length (red), predicted from obtained average elongation rate curves, together with the average HADA staining intensity at the cell pole after background correction (blue). Results are shown for a polar region defined to be within 0.51 $\mu$m (**A**) and 1.0 $\mu$m (**B**) of the cell tip.

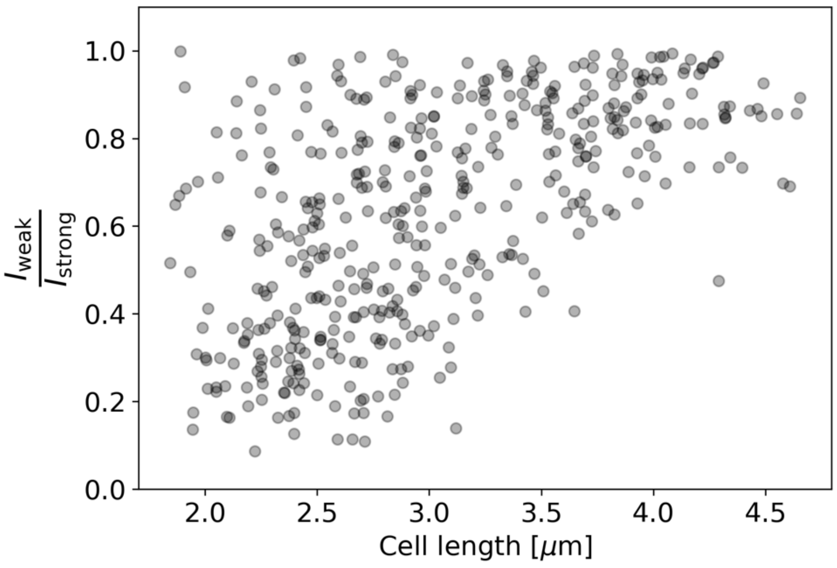


**Appendix 2-Figure 11 Ratio of intensities between the weaker and the stronger pole of each cell in the HADA staining experiment.** Polar intensities are calculated as described in Appendix 2-Figure 8. Here, $I_{\text{weak}}$ denotes the intensity of the cell pole with the weaker HADA intensity signal, and $I_{\text{strong}}$ denotes the intensity of the pole with the stronger signal. For NETOlike growth (Hannebelle et al., 2020), a clustering of values around 0 (before new end take off) and 1 (after new end take off) would be expected, which is not observed here.

#### Appendix 3

**Measurement noise estimate**

To obtain an estimate for the measurement noise from our time-series growth data, we make use of length measurements at subsequent time intervals. For short enough time intervals, the variance of the length differences between intervals can be used as a measure of the measurement noise. However, since we expect cellular growth to also significantly contribute to this variance within the 3-minute measurement interval, we have to separate out the two contributions.

To separate out the two contributions to the variance in subsequent length measurements, we write this variance as

$$\mathrm{Var}(l_m(t+\Delta t) - l_m(t)) \;=\; \mathrm{Var}(l(t+\Delta t) - l(t)) + 2\sigma_{\mathrm{n}}^2 \qquad \text{(A7)}$$

with $l_m(t)$ the measured length at time $t$, $l(t)$ the actual length at time $t$, and $\sigma_{\mathrm{n}}$ the standard deviation of the measurement noise. This expression can be derived by noting that for a single elongation trajectory, we have

$$l_m(t+\Delta t) - l_m(t) \;=\; l(t+\Delta t) + \xi - (l(t) + \xi) = l(t+\Delta t) - l(t) + \sqrt{2}\xi, \qquad \text{(A8)}$$

with $\xi$ the measurement noise. A solution for $\sigma_{\mathrm{n}}$ can be found if the functional form of $\mathrm{Var}(l(t), l(t+\Delta t))$ is known, by obtaining values for multiple $\Delta t$ and treating $\sigma_{\mathrm{n}}$ as a fitting parameter. To obtain this functional form, we make use of the observed linear growth regime after ~20 minutes (Main Text Figure 5). We observe that the elongation rate is approximately constant in this regime for cells of all birth lengths, and now assume that this is also true for cells individually within this regime. The contrary would imply that non-constant single-cell elongation rates precisely cancel out across time and birth lengths to produce linear growth, which seems biologically implausible.

For linearly growing single cells, the standard deviation of $l(t + \Delta t) - l(t))$ is proportional to $\Delta t$, implying that the term $\mathrm{Var}(l(t), l(t + \Delta t))$ is of the form

$$\mathrm{Var}(l(t + \Delta t) - l(t)) = c\Delta t^2 \,, \tag{A9}$$

with $c$ an unknown parameter. To simultaneously obtain $c$ and $\sigma_{\mathrm{n}}$, we fit Eq. (A7) under substitution of Eq. (A9) to the DivIVA labelled cell data over the regime between the onset of linear growth (18 min, black dashed line Appendix 3-Figure 1) and the first division event (36 min, grey dashed line Appendix 3-Figure 1). From this fit, we obtain the estimates $\sigma_n = 0.060 \pm 0.018$ μm and $c = 4.5x10^{-5} \pm 0.47 \mathrm{x} \mu\mathrm{m}^2 \mathrm{min}^{-2}$, where the error margins are determined via bootstrapping. This value of $\sigma_n$ is used in the correction procedure for assigned birth lengths described in Appendix 4.

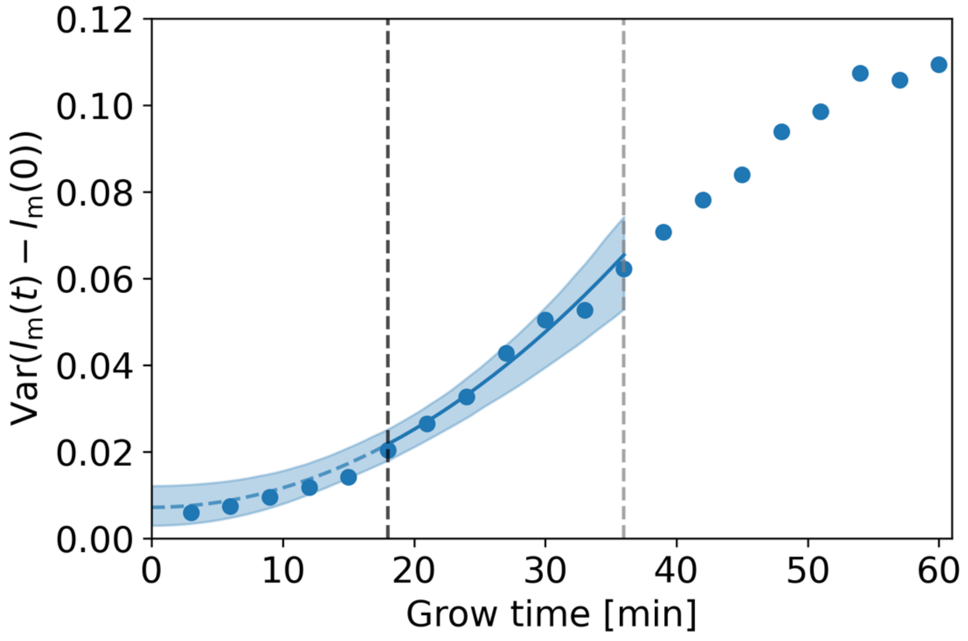

**Appendix 3-Figure 1 Estimation of measurement noise procedure.** Blue dots: variance of $l_m(t) - l_m(0)$ as a function of grow time for the DivIVA labelled cells, with $l_m(t)$ the measured cellular length at grow time $t$. A fit of Eq. (A7) under substitution of Eq. (A9) (blue line) is made to the points between the onset of linear growth (black dashed line) and the moment of first division (grey dashed line). The value of the extrapolated fit (blue dashed line) at t=0 is equal to $2\sigma_{\mathrm{n}}^2$, with $\sigma_{\mathrm{n}}$ the standard deviation of the measurement noise. The 95% confidence intervals of the model fit (blue shaded area) are obtained via bootstrapping.

#### Appendix 4

**Bias correction procedure for assigned birth lengths**

Before calculating average elongation rate curves, a statistical bias arising in the assignment of birth lengths to each curve needs to be corrected for. This bias is not specific to the inference method introduced in this paper, but arises in any procedure involving the assignment of lengths to a cells within a population, if there is noise in the measurement of individual cell lengths.

Due to measurement noise, cells will be assigned to birth lengths that systematically differ from their actual birth lengths. Specifically, given that the birth lengths in the population follow a symmetric, unimodal distribution, cells with a measured birth length larger than the population mean will on average be assigned a birth length that is larger than their actual length. Conversely, cells with a birth length smaller than the population mean will on average be assigned a birth length that is smaller.

The magnitude of the systematic deviation in the assignment of birth lengths is calculated as follows. Given that the cellular birth lengths follow a Gaussian distribution $P_l(l_b)$ with mean $\mu_l$ and standard deviation $\sigma_l$, and the measurement noise follows a Gaussian distribution $P_n(\Delta \mathrm{l})$ with mean 0 and standard deviation $\sigma_n$, the distribution of measured lengths will again be a Gaussian, with mean $\mu_m = \mu_l$ and standard deviation $\sigma_m = \sqrt{\sigma_l^2 + \sigma_n^2}$.

For a given measured birth length $l_m$, we now consider the probability distribution of corresponding actual birth lengths $P_l(l_b|l_m)$. This distribution is given by

$$P_l(l_b|l_m) = P_l(l_b)P_n(l_m - l_b). \tag{A4}$$

The product of two Gaussian distributions is again Gaussian, with a mean equal to

$$< l_b|l_m >= \frac{\sigma_n^2\mu_l + \sigma_l^2 \int l_b P_n(l_m - l_b)\mathrm{d}l_b}{\sigma_n^2 + \sigma_l^2} = \frac{\sigma_n^2\mu_l + \sigma_l^2 l_m}{\sigma_n^2 + \sigma_l^2}. \quad \text{(A5)}$$

Equation (A5) thus provides the transformation needed to remove the systematic bias in the assignment of birth lengths, and to determine the most likely birth length $l_b$ to a cell with a measured birth length $l_m$. For an estimation of the experimental measurement noise, see Appendix 3.

For the length increase since birth, there is no systematic bias once the bias in birth length has been removed. We can see this as follows. For each single-cell elongation trajectory, the measured length $l_m(t)$ at time $t$ is given by

$$l_m(t) = l_b + \Delta l_t + \xi, \quad \text{(A6)}$$

with $\xi$ the measurement noise and $\Delta l_t$ the length increase since birth at time $t$. As the measurement noise $\xi$ has a zero mean, there is no systematic bias in length increases after birth, provided that we have an unbiased estimate for the birth length $l_b$.

To test the derived correction procedure for assigned birth lengths, we performed a simulation of a population of growing cells, with the length measurement subject to noise. The measurement noise was sampled from a Gaussian, with the same standard deviation as estimated for experiment (Appendix 3). The single-cell growth mode was chosen as an input parameter. We analyzed two choices for input growth mode: linear (Appendix 4-Figure 1A, C, dashed lines) and exponential (Appendix 4-Figure 1B, D, dashed lines), with elongation rates comparable in magnitude to measured elongation rates.

For each single-cell growth mode, we applied our elongation rate inference procedure to simulated cell lengths subject to measurement noise. Without correcting for a bias in assigned birth lengths, we find a systematic deviation between inferred elongation rates and input elongation rates in both cases (Appendix 4-Figure 1A, B). With the implementation of the correction for assigned birth lengths, the input elongation rates are, however, accurately recovered (Appendix 4-Figure 1C, D).

Minor deviations from the input elongation rates can still be seen for exponentially growing cells (Appendix 4-Figure 1D), arising from applying a Gaussian smoothing to elongation curves that are locally nonlinear due to limited time resolution. However, this effect is small compared to the uncertainty on the inferred elongation rates.

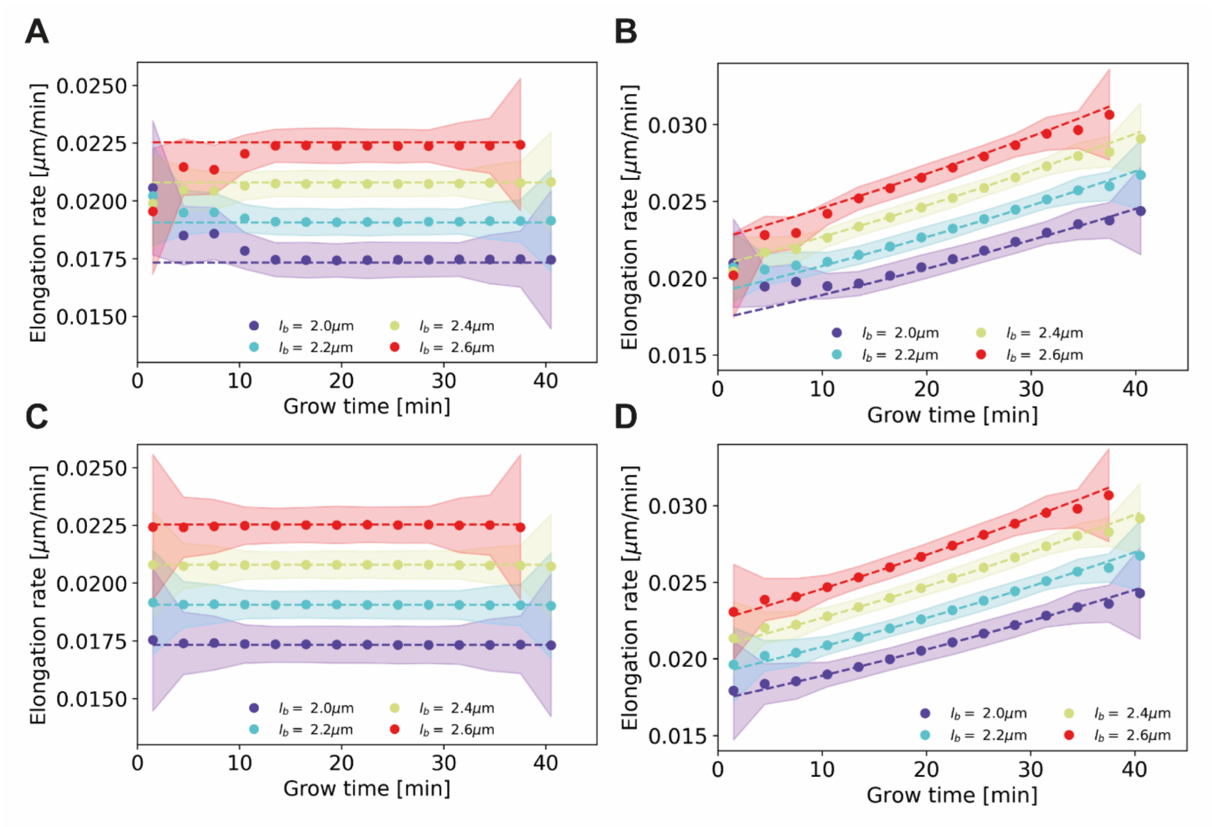


**Appendix 4-Figure 1 Elongation rate inference on simulated data sets, with and without bias correction procedure for assigned birth lengths.** For all panels: dashed lines: input elongation rates. Dots: mean inferred average elongation rates, obtained by applying our inference procedure to 1000 simulated data sets. Shaded areas: 2σ bounds on the inferred elongation rates. For all simulated data sets, the measurement noise is drawn from a Gaussian distribution with a standard deviation of 0.075 μm, matching the estimated experimental noise (Appendix 4). The population size and birth length distribution are chosen to match those observed for the DivIVA labelled cells. Simulation conditions: **(A)** Linear input elongation rates constructed by setting $l(t) = l_b + 0.26 l_b t$. No bias correction procedure for assigned birth lengths is applied. **(B)** Exponential input elongation rates constructed by setting $l(t) = l_b e^{0.26t}$. No bias correction procedure for assigned birth lengths is applied. **(C)** Input elongation rates as in (A). The bias correction procedure for assigned birth lengths is applied. **(D)** Input elongation rates as in (B). The bias correction procedure for assigned birth lengths is applied.

#### Appendix 5

##### Smoothing of elongation curves

We obtain elongation rate curves (Main Text Figure 5 and Figure 6 C) by taking a numerical derivative of smoothed growth trajectories. For the smoothing, a Gaussian smoothing procedure was used. In this procedure, a moving average is applied twice over groups of 3 subsequent time stamps of average elongation curves. As a check of the validity of the smoothing procedure, we also compare elongation rates before and after smoothing (Appendix 5-Figure 1).

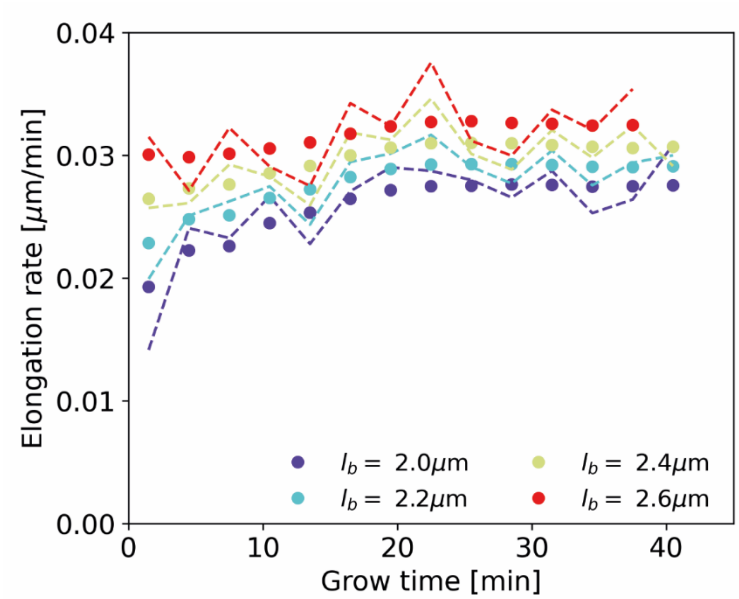


**Appendix 5-Figure 1** Average elongation rate curves obtained after Gaussian smoothing of the inferred average elongation curves (dots), together with average elongation rate curves obtained from unsmoothed average elongation curves (dashed lines).

#### Appendix 6

**Testing the elongation rate inference procedure**

To test our elongation rate inference procedure, we generated a simulated data set with elongation rates as inferred by our inference procedure for DivIVA labelled cells (Main Text Figure 5). The distribution of birth lengths and division lengths of the simulated cells are taken to match the experimentally observed distributions. On each simulated data point, a measurement noise as determined in Appendix 3 is applied. On the simulated data set subject to noise, we apply the assigned birth length correction procedure as described in Appendix 4, and subsequently apply our elongation rate inference procedure. We find that the input elongation rates are accurately recovered (Appendix 6-Figure 1), demonstrating the internal consistency of our inference approach.

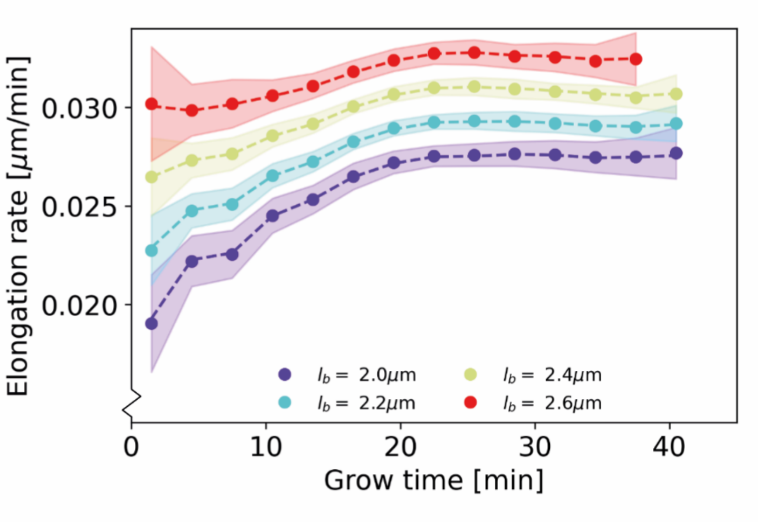


**Appendix 6-Figure 1 Recovery of inferred elongation rates from simulated growth**

Dashed lines: input elongation rates, as inferred for DivIVA labelled cells (Main Text Figure 5A). Dots: average of elongation rates inferred from simulated growth experiment. Shaded areas: 95% confidence intervals inferred from simulated growth experiment, obtained via bootstrapping.

#### Appendix 7

**Prediction of average elongation rate as a function of cell length**

To calculate the predicted average elongation rates shown in Main Text Figure 6C, we make use of our time-series data for wild-type cells, and the inferred mean elongation rates shown in Main Text Figure 5B.

We start by calculating the time-averaged elongation rate $\overline{l'}_i$ for each cell $i$ in the wild-type data set, where the prime denotes a time derivative, by dividing the length added between birth and division by the total growth time. We then assume that the elongation rate for a cell at a time $t$ is approximately given by a rescaling of the population-averaged elongation rates $L'(t, l_b)$ by the time-averaged elongation rate of the cell $\overline{l'}_i$. Specifically, we calculate the estimated elongation rate at time $t$ by

$$l'_i(t) = L'(t, l_b) \frac{\overline{l'}_i n_{\mathrm{i}}}{\sum_{t=0}^{t_{\mathrm{div}}^i} L'(t, l_b)}, \qquad \text{(A10)}$$

with $n_{\mathrm{i}}$ the number of time intervals in the growth trajectory of cell $i$, and $t_{\mathrm{div}}^i$ its division time. For times $t$ later than the first population division event $T_{\mathrm{div}}$, we obtain a value for $L'(t, l_b)$ by extrapolating the linear growth regime, setting $L'(t, l_b) = < L'(t, l_b) >_{20\ \mathrm{min}<t<T_{\mathrm{div}}}$.

From the ensemble $\{l'_i(t)\}$ of estimated elongation rates of all cells at each time since birth, we calculate the average elongation rate as a function of cell length by taking a moving average over the corresponding measured $\{l_i(t)\}$. The standard error on the mean is calculated from the standard deviation and the number of cells of each moving average bin.

### Appendix 8

**RAG Model fitting procedure**

The model fits shown in Main Text Figure 6 E-G are obtained via the ParametricNDSolve function in Mathematica. The obtained parameter values are shown in Tables 1 and 2.

**Appendix 8-Table 1 Parameter values obtained by fitting Main Text Eq. (2) to inferred elongation rate curves.** The values shown in column 4 and 6 are an average over the four birth lengths of each condition.

| Genotype | $l_b$ [μm] | $\beta\ [t^{-1}]$ | $<\beta>\ [t^{-1}]$ | $\frac{N(t=0)}{N^{\max}}$ | $<\frac{N(t=0)}{N^{\max}}>$ |
|---|---|---|---|---|---|
| *wild-type* | 2.1 | 0.088 | 0.085 | 0.67 | 0.62 |
| | 2.3 | 0.068 | | 0.62 | |
| | 2.5 | 0.093 | | 0.62 | |
| | 2.7 | 0.089 | | 0.58 | |
| *divIVA:divIVA-mCherry* | 2.0 | 0.109 | 0.088 | 0.69 | 0.80 |
| | 2.2 | 0.100 | | 0.77 | |
| | 2.4 | 0.087 | | 0.84 | |
| | 2.6 | 0.054 | | 0.88 | |
| *divIVA:divIVA-mCherry ΔrodA* | 1.7 | 0.063 | 0.087 | 0.61 | 0.64 |
| | 1.9 | 0.094 | | 0.65 | |
| | 2.1 | 0.084 | | 0.64 | |
| | 2.3 | 0.11 | | 0.65 | |

**Appendix 8-Table 2 Parameter values obtained by fitting Main Text Eq. (3) to inferred elongation rate curves.** The values shown in column 5 and 7 are an average over the four birth lengths of each condition.

| Genotype | $l_b$ [μm] | $\beta\ [t^{-1}]$ | $\gamma\ [t^{-1}]$ | $< \beta e^{\gamma t} >_{t<20min}\ [t^{-1}]$ | $\frac{N(t=0)}{N^{\max}}$ | $< \frac{N(t=0)}{N^{\max}} >$ |
|---|---|---|---|---|---|---|
| *wild-type* | 2.1 | 0.016 | 0.162 | 0.13 | 0.72 | 0.67 |
| | 2.3 | 0.039 | 0.086 | | 0.67 | |
| | 2.5 | 0.058 | 0.080 | | 0.65 | |
| | 2.7 | 0.082 | 0.050 | | 0.62 | |
| *divIVA:divIVA-mCherry* | 2.0 | 0.072 | 0.06 | 0.14 | 0.71 | 0.82 |
| | 2.2 | 0.050 | 0.09 | | 0.79 | |
| | 2.4 | 0.025 | 0.14 | | 0.86 | |
| | 2.6 | 0.005 | 0.25 | | 0.92 | |
| *divIVA:divIVA-mCherry ΔrodA* | 1.7 | 0.023 | 0.094 | 0.12 | 0.67 | 0.68 |
| | 1.9 | 0.039 | 0.092 | | 0.69 | |
| | 2.1 | 0.064 | 0.050 | | 0.67 | |
| | 2.3 | 0.064 | 0.100 | | 0.68 | |

#### Appendix 9

**Population simulation method**

The goal of the population growth simulations is to obtain the distribution of cellular birth lengths assuming two different growth modes: asymptotically linear and exponential elongation. Both simulations extract all necessary growth parameters and distributions from the experimental data. For the asymptotically linear growth mode, the simulation serves as a check whether the assumed growth mode indeed recovers the correct cellular length distribution. For the exponential growth scenario, the simulation reveals the cellular length distribution an exponential grower would have if it had inherent noise levels similar to *C. glutamicum* allowing for a fair comparison. Both simulations start with a single cell and continue for 20 generations, after which the birth lengths of the last generation are binned and plotted. Repeated simulations with different lengths of the starting cell do not show discernable differences.

<u>Exponential growers</u>

For the exponential growers, cells are assumed to elongate according to

$$l(t) = l_b \exp(\alpha t) + \zeta(t) \qquad \text{(A11)}$$

The exponential growth rate $\alpha$ is chosen as the slope of the linear fit of $\ln\left(\frac{l_d}{l_b}\right)$ versus $t_d$ that intersects the origin, as shown in Main Text Figure 3B. A size-additive noise term is indicated by $\zeta(t)$, which will be specified below at the time of division. For a cell with a given birth length $l_b$, the target final length $l_t$ is determined via a linear fit of $l_b$ versus $l_d$, as shown in Main Text Figure 3A. The target growth time $t_t$ is then given by $t_t = \frac{1}{\alpha}\ln\left(\frac{l_t}{l_b}\right)$. A time-additive noise term $\Delta t$ is added to $t_t$ according to experimentally observed growth time variations (Appendix 9-Figure 1D). Additionally, a size-additive noise term $\Delta l$ encodes the division length variation due to $\zeta(t)$, which is also directly obtained from experiment (Appendix 9-Figure 1E).

The full expression for the division length $l_d$ is then given by

$$l_d = l_b \exp(\alpha(t_t + \Delta t)) + \Delta \mathrm{l} \quad \text{(A12)}$$

At division, the characteristic v-snap of *C. glutamicum* occurs, separating the two daughter cells. During this v-snap, the length of the daughter cells rapidly increases: the average measured birth length is 0.57 times the average measured division length (2.3 $\mu$m and 4.0 $\mu$m respectively), instead of the expected ratio of 0.5. To account for this v-snap effect, we calculate the distribution of added lengths during the v-snap. We find that the average added length depends on the division length: longer cells add less length during the v-snap than shorter cells (Appendix 9-Figure 1B). To take this length dependence into account, we subdivide the data set into three division length bins, and obtain a distribution of added lengths during the v-snap for each bin. When a simulated cell divides, an added length during v-snap is randomly drawn from the distribution corresponding to its division length.

After division, the length asymmetry of the two daughter cells is chosen by drawing a random value from the experimentally observed division asymmetry distribution (Appendix 9-Figure 1C) corresponding to the obtained division length. This distribution is found to be narrower for the shortest birth lengths (Appendix 9-Figure 1C), thus two distributions are used.

<u>Asymptotically linear growers</u>

For asymptotically linear growers, cells are assumed to elongate according to

$$l(t) = l_b + \lambda t + \gamma(\exp(-\beta t) - 1) + \eta(t), \quad \text{(A13)}$$

which is obtained by inserting Main Text Eq. (3) into Main Text Eq. (1), integrating and grouping constant terms into $\lambda$ and $\gamma$. An additive noise term $\eta(t)$ is added to this to account for single-cell variability around the inferred average growth trajectory. We assume the cells to have the same target final length $l_t$ as in the exponentially growing scenario, determined via a linear fit of $l_b$ versus $l_d$. For cells close to observed division times, the term proportional to $\gamma$ can be approximated as being constant in time, simplifying the growth mode to linear growth (Main Text Figure 5A). A time-additive noise term $\Delta t$ will then act as size-additive noise and can thus be absorbed into one additive noise term $\Delta l$, obtained from the experimental distribution of final sizes $l_d$ around the target final sizes (Appendix 9-Figure 1F). The expression for the division length is thus given by

$$l_d = l_t + \Delta l. \tag{A14}$$

The division asymmetry and v-snap effect are incorporated in the same way as for the exponential grower simulation.

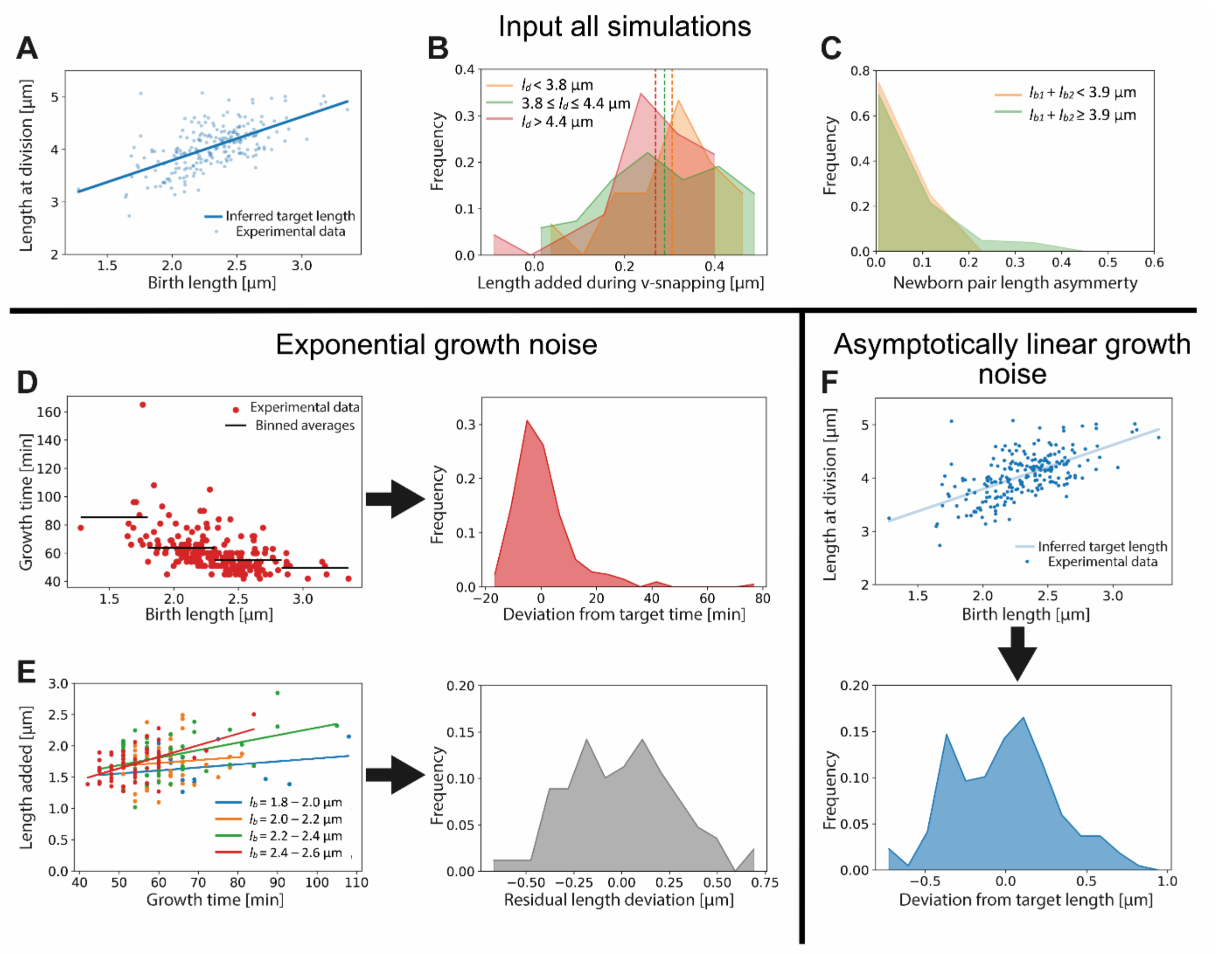


**Appendix 9-Figure 1** Input used for simulations of exponential and asymptotically linear growth. For both simulations, a linear fit of the division length versus birth length is used to define a target length (**A**). The length added during the v-snap at division is randomly drawn from the distribution corresponding to the division length of the simulated cell (**B**). The experimental data is divided into three subpopulations according to division length (red, green and orange distributions), as the average length added during v-snap decreases with division length (dashed lines). The asymmetry of the daughter cells is randomly drawn from the distribution corresponding to the combined length of the simulated daughters (**C**). As the asymmetry is lower for the smallest daughter cells, the experimental data is divided into two subpopulations (red and green distributions). For the simulation of exponential growth, two noise sources are needed as input. The time-additive noise is randomly drawn from the distribution of deviations from target growth times (**D**). This distribution is obtained from the deviations of single-cell growth times from the average of their birth length bin. All growth variability not captured by growth time variations is calculated for four narrow birth length bins (blue, orange, green and red points) (**E**). From the distribution of deviations of added lengths from a linear fit for each initial size bin, a size-additive noise term is randomly drawn. For the linear growth simulation, only a single additive noise term is required, which is randomly drawn from the distribution of deviations of cells lengths at division from the target division length (**F**).

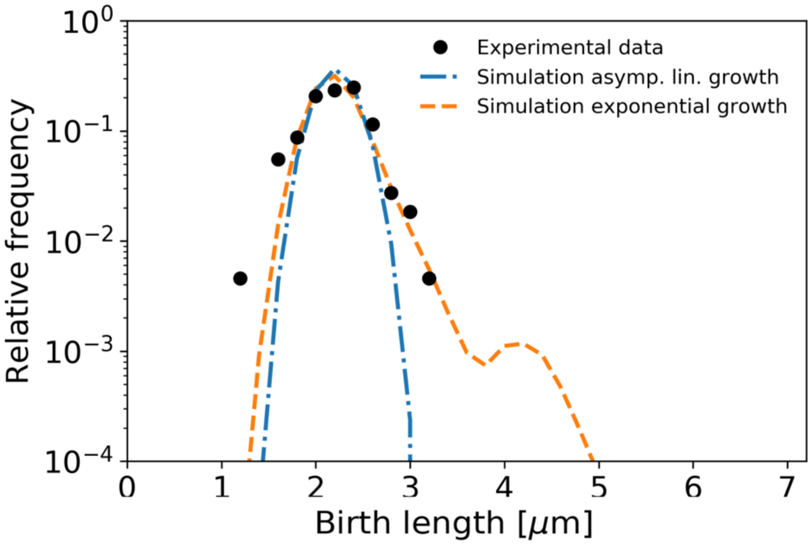


**Appendix 9-Figure 2** Birth length distributions as in Main Text Figure 7, but with single-cell variability in division symmetry, growth time, and (residual) length deviation reduced by a factor 3. The second peak in the length distribution of exponential growth is attributed to the large time deviation of one single cell seen in Appendix 9-Figure 1D.
